# Trehalose Transport Dynamics Underpin a Metabolic Trade-Off between Exogenous Uptake and Endogenous Synthesis in Lepidopteran Insects

**DOI:** 10.1101/2025.09.11.675667

**Authors:** Bhagyashri Y. Chaudhari, Vikram J. Nichit, Vitthal T. Barvkar, Rakesh Shamsunder Joshi

## Abstract

Trehalose is the major insect hemolymph sugar and plays a diverse role. Its level is regulated endogenously by the dynamics of biosynthesis and distribution by sugar transporters (STs). The metabolic trade-off between trehalose synthesis and uptake remains poorly understood, despite its critical role in homeostasis. Here, we examined the role of a gut-specific trehalose transporter, *HaST46*, in regulating this metabolic trade-off in *Helicoverpa armigera*, a Lepidopteran pest model. Integrated transcriptomics analysis and functional analyses revealed that *HaST46* acts as a diet-responsive transporter, localised to the posterior midgut, with trehalose preference. Its expression is modulated in response to dietary trehalose availability, enhancing the efficient exogenous trehalose uptake while attenuating its endogenous synthesis and conserving energy. Functional perturbation through overexpression and silencing revealed a feedback-regulated mechanism in which *HaST46* expression showed strong correlation with trehalose metabolising enzymes and other HaSTs isoforms to maintain systemic trehalose homeostasis. Overall, our findings reveal a metabolic trade-off between exogenous trehalose uptake and endogenous synthesis mediated by gut-specific sugar transporters.

## 1. Introduction

Insects exhibit a remarkable metabolic adaptation that contributes to their ecological success. Carbohydrates serve as a primary energy source for fuelling flight, development, reproduction, diapause, and stress tolerance (Thorat et al. 2012; Feofilova et al. 2014). Under stress conditions, insects must precisely regulate their energy acquisition, storage, and mobilisation (Zhang et al. 2019). Trehalose is the predominant hemolymph sugar in insects, fulfilling both structural and metabolic functions (Becker et al. 1996; Matsushita and Nishimura 2020;,Nishimura 2020;,Murphy and Wyatt 1965;,Shukla et al. 2015). Its transport and metabolism are essential for insect growth and adaptability (Tellis, Kotkar, et al. 2023). Trehalose is primarily synthesised in the fat body through a conserved pathway involving *trehalose-6-phosphate synthase* (*TPS*) and *trehalose-6-phosphate phosphatase* (*TPP*) (Murphy and Wyatt 1965; Becker et al. 1996). Following synthesis, trehalose is transported into the hemolymph and then distributed to the target tissues as per the energy demand through trehalose transporters (TRETs) (Tellis, Chaudhari, et al. 2023).

Insects can ingest a wide range of foods to obtain complementary nutrients during nutritional scarcity (Koyama et al. 2020;,Raubenheimer and Jones 2006). They further optimise energy utilization through metabolic balancing, enabling adaptation to diverse dietary conditions (Smith et al. 2018;,Chown and Gaston 1999). Trehalose synthesis requires a continuous supply of ATP and carbon skeletons from glycolysis, glycogenolysis, or gluconeogenesis (Miyamoto and Amrein 2017). Depending on resource availability, resource insects may shift their metabolism to suppress endogenous synthesis and prioritise exogenous uptake (Izadi 2025). Hemolymph trehalose levels range from 5 to 50mM, which is maintained by continuous exchange via TRETs for effective utilisation (Thompson 2003;,Al Hinai et al. 2025;,Wyatt 1967; Kikawada et al. 2007). TRETs are key in regulating metabolic trade-offs under controlled and stressed conditions (Leyria et al. 2020). The midgut is the primary site of sugar and nutrient absorption, detoxification, and enzymatic activity (Huang et al. 2015). To meet the metabolic demand, insects rely on sugar transporters such as TRETs, sodium-glucose co-transporters (SGLTs), and facilitative glucose transporters (GLUTs) to manage dietary sugar acquisition and use (Chaudhari et al. 2025).

Phytophagous insects such as *H. armigera* can acquire trehalose directly from their diet (Shukla et al. 2015; Tellis, Kotkar, et al. 2023). In *H. armigera*, trehalose-specific sugar transporters (*HaSTs*) showed diversity and spatio-temporal expression patterns (Tellis, Chaudhari, et al. 2023). In dipteran and hymenopteran insects, glucose absorption is primarily mediated by SGLTs. and GLUTs (Wang et al. 2016). In *Drosophila melanogaster*, the diet-responsive GLUT homolog CG4797 and SGLT homolog dSGLT1 have been shown to influence development, fertility, and survival (Chaudhari et al. 2025). The activity of these transporters is closely linked to the insulin and TOR signalling pathways, facilitating metabolic plasticity by coupling nutrient availability with growth regulation (Riera et al. 2016). This critical trait allows insects to cope with environmental stressors, shift between trophic modes, and optimise energy expenditure during metamorphosis or overwintering (Danks 2007). The interplay between trehalose metabolism and other physiological systems, including neuroendocrine signalling, symbiotic relationships, and hormonal regulation, facilitates nutrient sensing, energy homeostasis, and ecological adaptability (Tellis, Kotkar, et al. 2023).

Despite advances in understanding trehalose metabolism within insects, the integrated dynamics between endogenous trehalose synthesis and exogenous uptake remain poorly understood in many model and non-model organisms. Lepidopteran pests such as *H. armigera* serve as valuable models for metabolic studies due to their high dietary adaptability, rapid development, and stress resilience (Riaz et al. 2021). Elucidating how these insects regulate trehalose metabolism in response to diet offers insights into the metabolic homeostasis and its impact on physiological processes (Dawkar et al. 2016;,Guan et al. 2024;, Tian et al. 2025;, Zhou et al. 2023). Here, we shed light on the functional role of the trehalose transporter, *HaST46*, in mediating the metabolic trade-off between trehalose’s exogenous uptake and endogenous synthesis. Through differential gene expression analysis, qRT-PCR, LC-MS-based metabolomics, enzyme activity assays, and genetic manipulation, we investigated how dietary trehalose influences *HaSTs* dynamics and activity, mainly in the gut. This study advances the knowledge of sugar transport, metabolic homeostasis and nutritional physiology.

## 2. Results

### 2.1 *HaST46* showed posterior midgut-specific expression at the transcript and protein levels

We analyzed the gut-specific expression and proteomic presence of selected putative trehalose transporters, *HaSTs*, using publicly available *H. armigera* transcriptomic and proteomic datasets (Ioannidis et al. 2022). Our previous study elucidated five putative trehalose transporters with selective abundance in specific tissues, among which *HaST46* and *HaST69* were found to be present in gut-specific transcriptomic data **(Figure 1a)**. While only *HaST46* showed expression in proteomic data, indicating its proteomic level abundance in the mid-gut **(Figure 1b)**. In the transcriptomics dataset, *HaST46* and *HaST69* showed high expression levels across larval stages (L2, L3, L4) and in different gut regions, including foregut (FG), anterior midgut (AMG), middle midgut (MMG), posterior midgut (PMG), and hindgut (HG) in fifth instar (L5) larvae. *HaST46* expression peaked in the gut during the L4 larval stage and in the PMG of L5 larvae **(Figure 1a)**. Given that feeding activity peaks during the fourth and fifth larval instars, elevated expression of the sugar transporter likely supports increased metabolic demand. Collectively, these data highlight that *HaST46* is a putative key facilitator of sugar transport across the PMG during the energetically demanding developmental stages and the period of active feeding. Transcriptomics comparisons were conducted between the gut compartments of L5 larvae fed an artificial diet (AD) versus those fed a plant diet (PD)-cotton to assess the diet-dependent expression dynamics. Interestingly, *HaST46* expression was elevated exclusively in the PMG of L5 larvae fed a plant diet (PD) compared to AD, whereas *HaST69* exhibited the opposite trend **(Figure 1c-d)**. The presence of HaST46 in the PMG, which is a primary site for nutrient absorption during active feeding, suggests its putative role in sugar absorption from feed. The selective higher expression of *HaST46* in larvae consuming a plant diet supports its involvement in sugar assimilation from plant (complex) food materials. This further suggests that the dietary composition possibly compensates for the metabolic demand.

**Figure 1.**
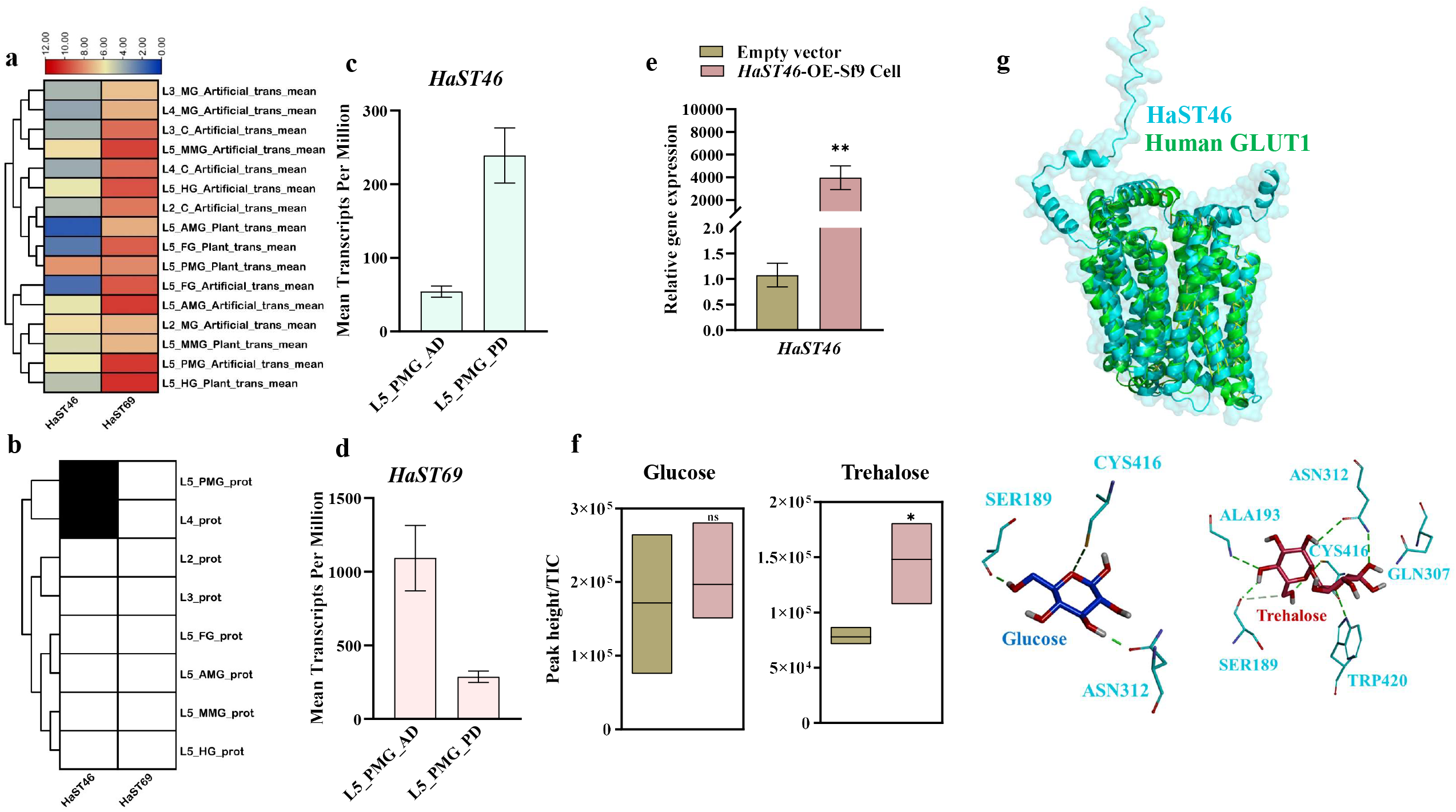
Spatial expression and functional characterisation of *HaST46*. (a) Protein abundance across larval stages (L2, L3, L4) and gut regions (FG: foregut; AMG: anterior midgut; MMG: median midgut; PMG: posterior midgut; HG: hindgut) in L5 larvae reared on an artificial diet. (b) DGE patterns comparing larvae fed on an artificial diet and a plant-based diet (cotton) across different gut compartments. (c, d) Expression levels of *HaST46* and *HaST69* in the posterior midgut under the two dietary regimes. Data represent mean ± SEM of biological replicates. Statistical significance was determined using an unpaired *t*-test (*P* < 0.05). (e) Verification of *HaST46* overexpression by qRT-PCR. (f) Quantification of D-glucose and trehalose in cell supernatants and lysates using LC–MS/MS after supplementation of the culture medium with glucose (5 mM, 10 mM) or trehalose (5mM, 10mM) at 0, 5, and 10 min intervals. All data are presented as mean ± SEM. Statistical analysis was performed using an unpaired *t*-test (*P* < 0.05). (g) Structural conservation of HaST46 with human GLUT1 transporter. Molecular interaction of HaST46 with glucose and trehalose.

### 2.2. HaST46 showed a preference toward trehalose transport

Functional assays with 10mM trehalose exposure to Sf9 cells overexpressing HaST46 revealed a significant increase in intracellular sugar accumulation in *HaST46*-expressing cells compared with the vector control **(Figure 1e and f)**. LC-MS based quantification revealed that *HaST46*-expressing cells accumulated ∼1.5-fold mor trehalose relative to controls, while glucose uptake exhibited a moderate increase (∼1.2-fold), suggesting a trehalose preference of HaST46. Furthermore, the structural alignment of HaST46 with the human GLUT1 transporter (PDB: 4PYP) showed a high degree of conservation in transmembrane domain architecture **(Figure 1g)**. Molecular docking studies further revealed that CYS416 and ASN312 of HaST46 formed conserved hydrogen bonds with both glucose and trehalose. At the same time, TRP420 and ALA193 contributed additional interactions that stabilized trehalose binding **(Figure 1g)**. These findings support the hypothesis that HaST46 functions as a trehalose-preferred transporter, likely contributing to sugar uptake and homeostasis in *H. armigera*.

### 2.3. Exogenous trehalose feeding upregulates *HaST46* expression and trehalose catabolism genes in *H. armigera*

Upon exogenous trehalose feeding, we observed optimal growth and fitness in the case of 50mM trehalose containing diet. In comparison to this, feeding with 10mM trehalose doesn’t showed any noteworthy physiological impact, and 100 mM trehalose lead to sugar-associated toxicity **(Supplementary File S5, Figures S1 and S2)**. Feeding of 50mM trehalose induced significant changes in the expression of trehalose transporters (TRETs) and metabolic genes as revealed by qRT-PCR analysis. *HaST46* expression was upregulated upon trehalose feeding **(Figure 2a)**. In response to this, *HaTPS/TPP* were downregulated, while *HaTreh-1* showed upregulation upon feeding a diet with 50mM trehalose **(Figure 2b-c)**. This alteration in the *HaTPS/TPP* expression level suggests a putative reduction in the energy-intensive endogenous trehalose biosynthesis and increased hydrolysis of exogenous trehalose for energy needs. Notably, *HaST69* was downregulated on a 50mM TD, indicating that it is potentially not involved in trehalose absorption from the gut **(Supplementary File S5, Figure S3)**. Residual TPP activity was slightly reduced, while trehalase activity was high in larvae fed the 50mM TD (p < 0.01) **(Figure 2d-e)**. Metabolomic quantification showed that the insect fed on the trehalose-containing diet showed relatively more accumulation compared to the control **(Figure 2f)**. Furthermore, increased accumulation of ATP and low ADP/AMP indicates little demand for ATP hydrolysis in trehalose fed insects **(Figure 2f)**.This evidence suggests a preference for hydrolytic metabolism under elevated dietary trehalose, which is likely to convert absorbed trehalose into glucose for systemic energy needs.

**Figure 2.**
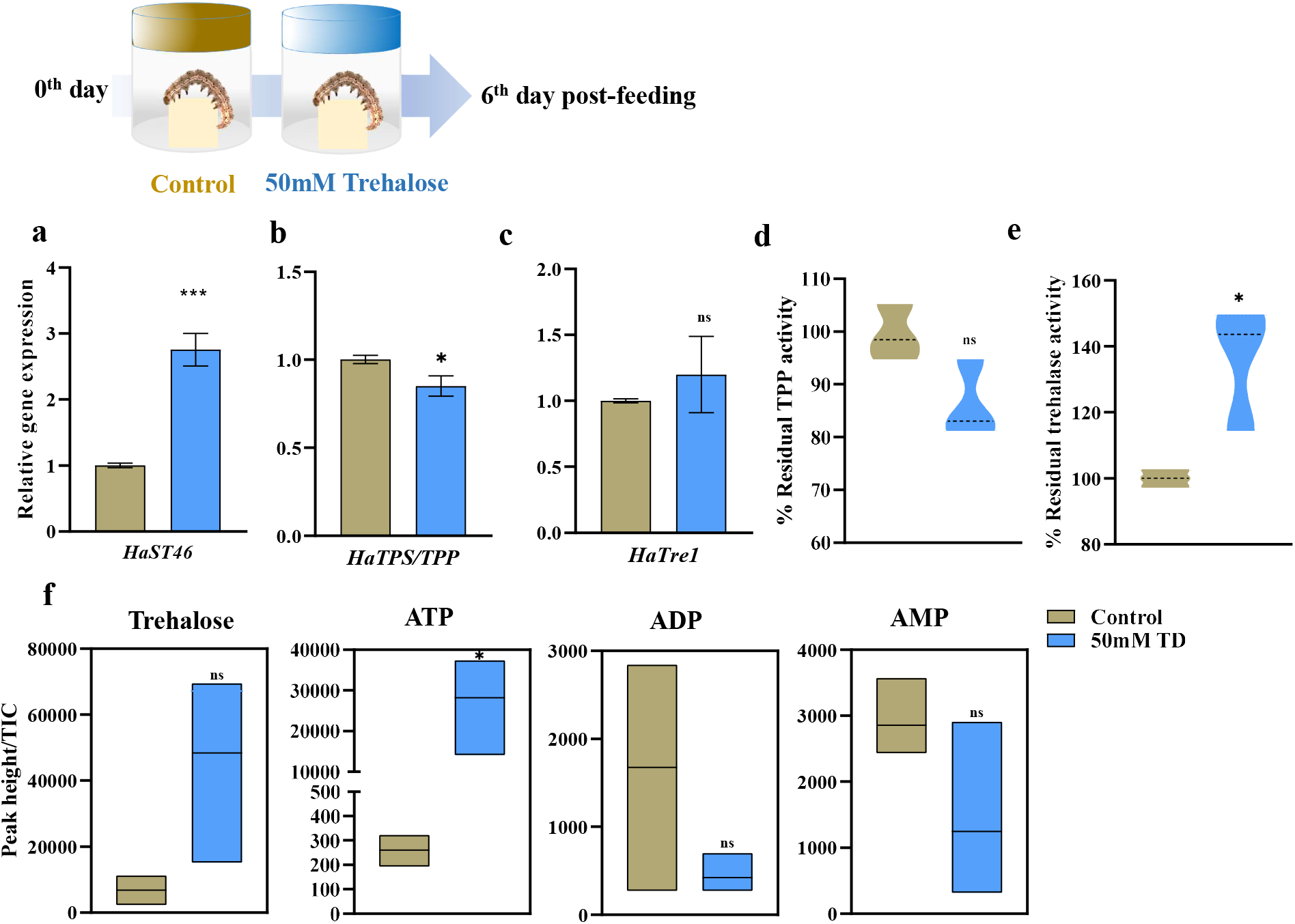
Gene expression and enzymatic activity upon exogenous trehalose feeding in *H. armigera*. (a–c) qRT-PCR analysis of *HaST46* and trehalose metabolism-related genes following dietary trehalose supplementation. Transcript abundance was normalised to reference genes and expressed as fold change (2^^−ΔΔCt^). (d, e) In vivo activity of trehalase and trehalose-6-phosphate phosphatase (TPP) in larvae fed a 50mM trehalose-enriched diet or an artificial diet. (f) Quantification of trehalose, ATP, ADP and, AMP in whole larvae using LC–MS under the same dietary conditions. Data are presented as mean ± SEM, with statistical significance assessed by an unpaired *t*-test (*P* < 0.05).

### 2.4. *HaST46* showed gut-specific upregulation in response to trehalose feeding

Tissue-specific gene expression upon 50mM trehalose feeding was further analyzed to elucidate the dynamics of trehalose transport. The relative expression of *HaST46* and *HaTPP* was reduced in hemolymph **(Figure 3a-b)**, while the expression of *HaTreh1* and *HaTreh2* was nearly unchanged **(Figure 3c-d)**. The gut showed a significant ∼3-fold increase in *HaST46* expression under a 50mM TD **(Figure 3e)**, indicating an increase in the need to transport and use excess trehalose in the gut tissue. As the midgut is the primary site for transporter-mediated dietary trehalose uptake, *HaST46* overexpression was found to be relevant. *HaTPS/TPP* expression was found to be unchanged, while *HaTreh1* and *HaTreh2* expression were significantly high **(Figure 3f-h)**. *HaST46* expression in the fat body (key tissue for trehalose synthesis) was found to be unchanged **(Figure 3i)**. Interestingly, the expression of *HaTPS/TPP* decreased significantly, suggesting a reduced endogenous trehalose biosynthesis **(Figure 3j)**. *HaTreh1* and *HaTreh2* expression were relatively high in the fat body of trehalose-fed insects **(Figure 3k-l)**. These observation suggests that dietary trehalose downregulates internal trehalose synthesis through feedback inhibition.

**Figure 3.**
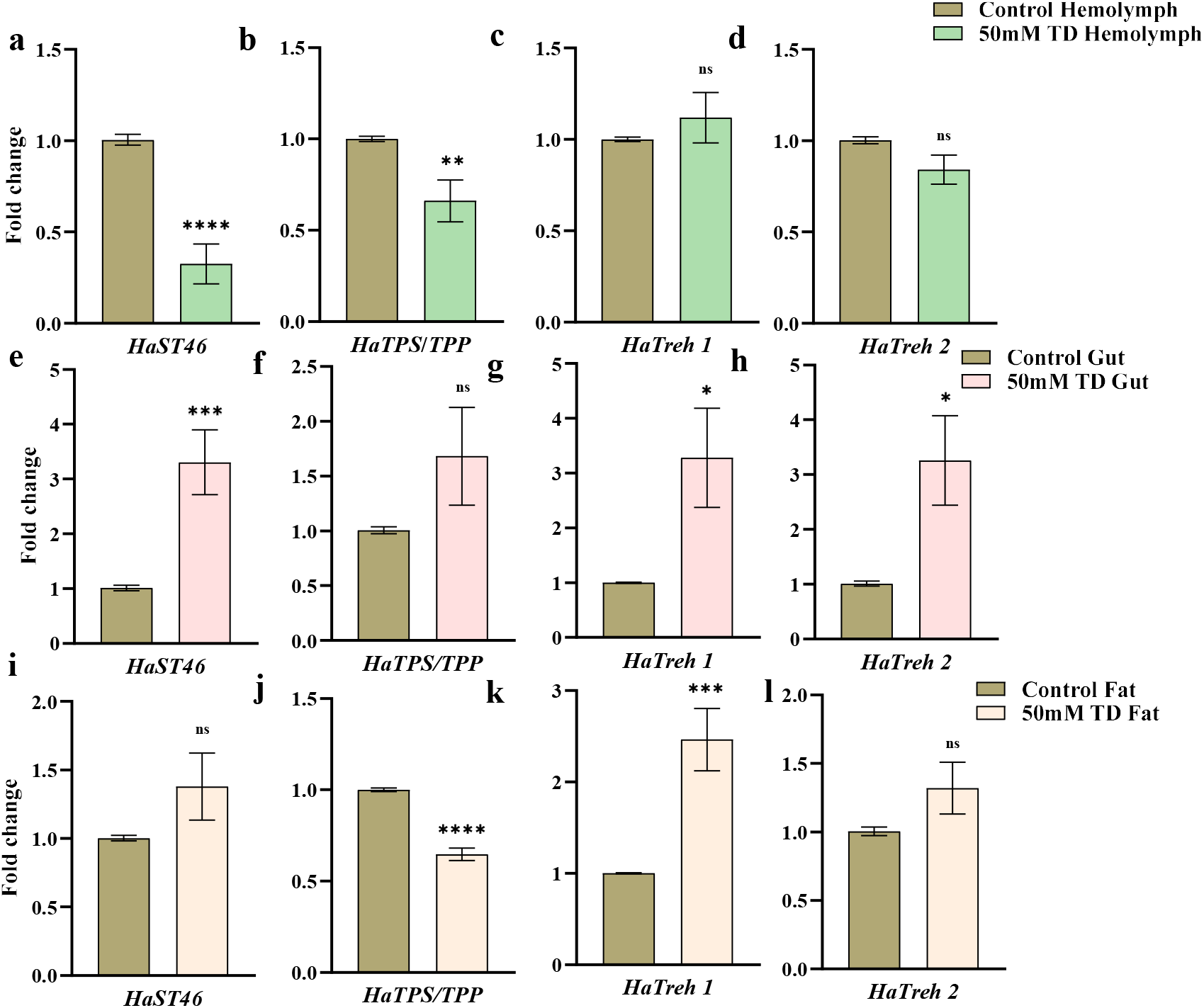
Tissue-specific expression of *HaST46* and trehalose metabolism genes following trehalose feeding. (a–c) qRT-PCR analysis showing expression patterns in haemolymph, fat body, and gut tissues. Transcript levels were normalised to reference genes and expressed as fold change (2^^−ΔΔCt^) relative to control. Data represent mean ± SEM, with statistical differences determined by an unpaired *t*-test (*P* < 0.05).

### 2.5. Metabolomic Profiling Reveals Trehalose Trade-off

LC-MS analysis was performed for hemolymph of insects fed on artificial diet (AD) and a trehalose-rich diet (TD) to quantify systemic metabolic changes. TD-fed larvae exhibited relatively elevated levels of trehalose, particularly in the hemolymph **(Supplementary File_S5, Figure S5)**. In addition, downstream glycolytic intermediates such as glucose-6-phosphate and pyruvate were elevated, supporting the notion that absorbed trehalose is rapidly catabolized into usable energy. These results confirm that trehalose supplementation alters gene expression and shifts the metabolic landscape toward catabolic flux, optimizing energy production in the presence of abundant dietary sugar in *H. armigera*.

### 2.6. *HaST46* knockdown induces compensatory upregulation of the trehalose anabolism TPS*/TPP* gene

RNA interference (RNAi) was employed to silence gut-prevalent *HaST46* expression. We achieved ∼80% reduction in the *HaST46* transcripts levels compared with the empty vector **(Figure 4a)**. Notably, insects with *HaST46* silencing that were fed a 50 mM TD exhibited a compensatory increase in *HaST46* expression, suggesting a feedback mechanism and highlighting its critical role in trehalose absorption from the gut **(Figure 4b)**. Furthermore, trehalase activity was elevated in TD-fed insects compared with AD-fed insects **(Figure 4c-d)**. LC-MS analysis revealed a decrease in trehalose levels in the insect body upon silencing *HaST46*, indicating putative involvement in the dietary trehalose absorption and resulting in an overall decrease in endogenous trehalose content **(Figure. 4e)**. However, upon trehalose feeding to *dsHaST46* insects, the trehalose level was almost the same as in the control **(Figure 4f)**. These findings highlight the potential role of *HaST46 in* trehalose transport. The impact of *HaST46* silencing on nutritional efficiency was assessed by measuring nutritional indices in both the control and silenced groups. Nutritional indices were almost the same in both groups, such as the control and *HaST46* silenced insects **(Supplementary File_S5, Figure S8-9)**.

**Figure 4.**
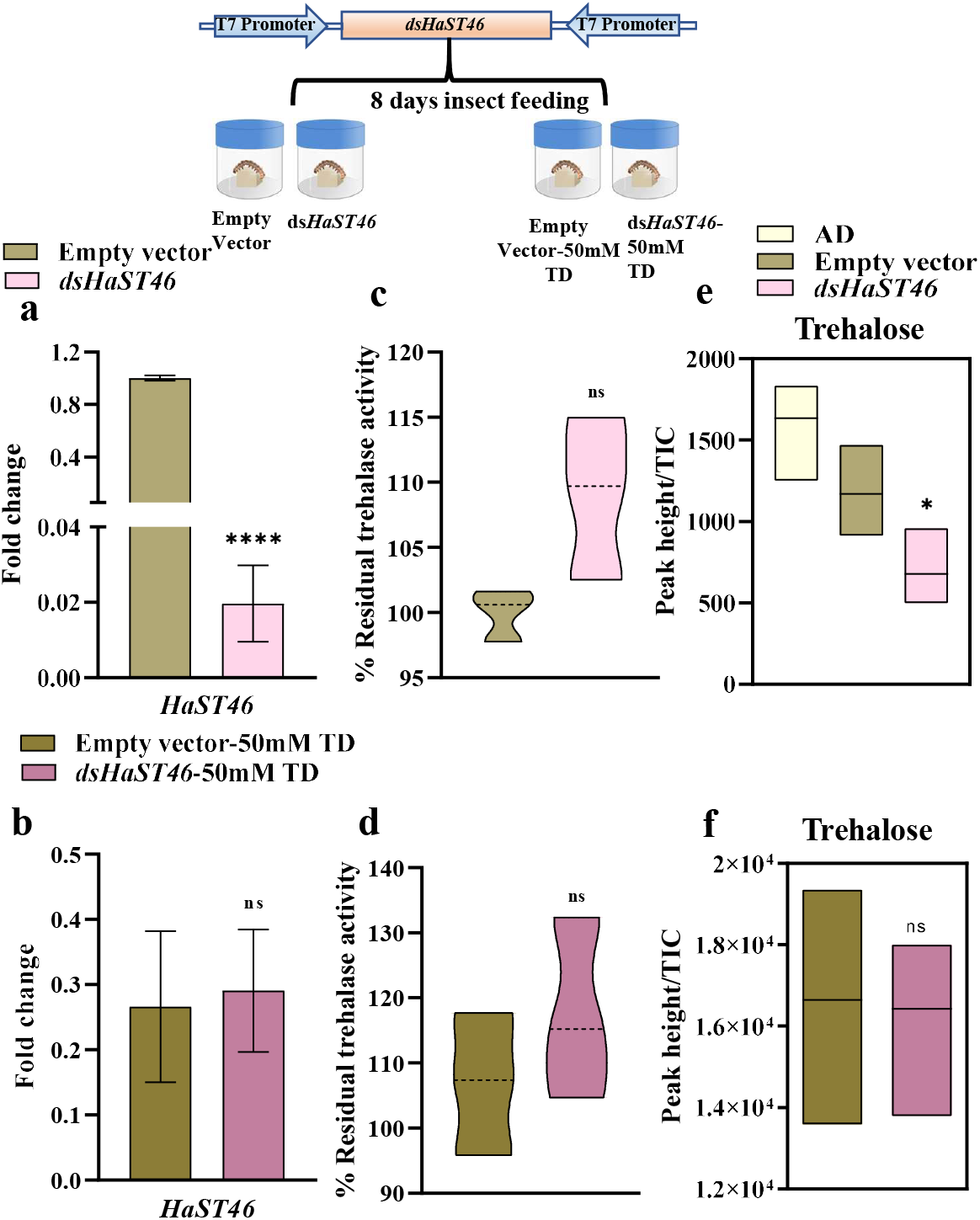
Silencing of *HaST46* in *H. armigera*. (a, b) qRT-PCR analysis of *HaST46* expression in whole-body samples from control insects (empty L4440 vector) and dsRNA-treated insects, either fed on an artificial diet or an artificial diet supplemented with 50mM trehalose. (c, d) In vivo trehalase and TPP activities under the same conditions. (e, f) Trehalose levels quantified in whole insects using LC–MS/MS. All values are mean ± SEM, with statistical analysis performed using an unpaired *t*-test (*P* < 0.05).

The expression of other trehalose transporter genes (*HaST09, HaST29, HaST64*, and *HaST69*) was examined in *dsHaST46*-fed larvae to explore potential compensatory mechanisms. Silencing *HaST46* resulted in significant downregulation of these genes, suggesting overall impact due to reduced trehalose content upon silencing (Supplementary File_S5, Figure 12a). *HaST09, HaST29, HaST64*, and *HaST69* exhibited upregulation in exogenous trehalose, although the functional significance of these changes requires further investigation (Supplementary File_S5, Figure 12b). These results underscore the pivotal role of *HaST46* in modulating trehalose transport and metabolism.

### 2.7. *HaST46* overexpression showed gut specificity and may enhance dietary trehalose

Transient overexpression of *HaST46* was confirmed with qRT-PCR analysis, showing ∼1000-fold higher expression of *HaST46* in treated larvae compared with the control **(Figure 5a)**. *HaST46* overexpression in larvae did not significantly alter trehalase activity, suggesting a maintained balance between trehalose synthesis and absorption to preserve homeostasis **(Figure 5b)**. Further, we observed the upregulation of *HaST46* in all three tissues of the *HaST46-OE* insects upon trehalose feeding compared with the control group. Interestingly, there was a significant upregulation of *HaST46* in the gut, indicating the potential preferred localization of overexpressed *HaST46* in the gut tissue **(Figure 5c-e)**. However, there was a significant increase in the *HaST46* transcript level in a group of insects fed on a trehalose-rich diet in the gut, followed by hemolymph and fat **(Figure 5c-e)**. Trehalose levels were slightly high in TD-fed *HaST46*_OE insects **(Figure 5f-g)**. At the same time, trehalase activity was also found to be high in TD-fed *HaST46*_OE insects **(Figure 5h)**. These results suggest that *HaST46* facilitates the uptake of dietary trehalose and reinforces its role in sugar homeostasis and energy balance in Lepidopterans.

**Figure 5.**
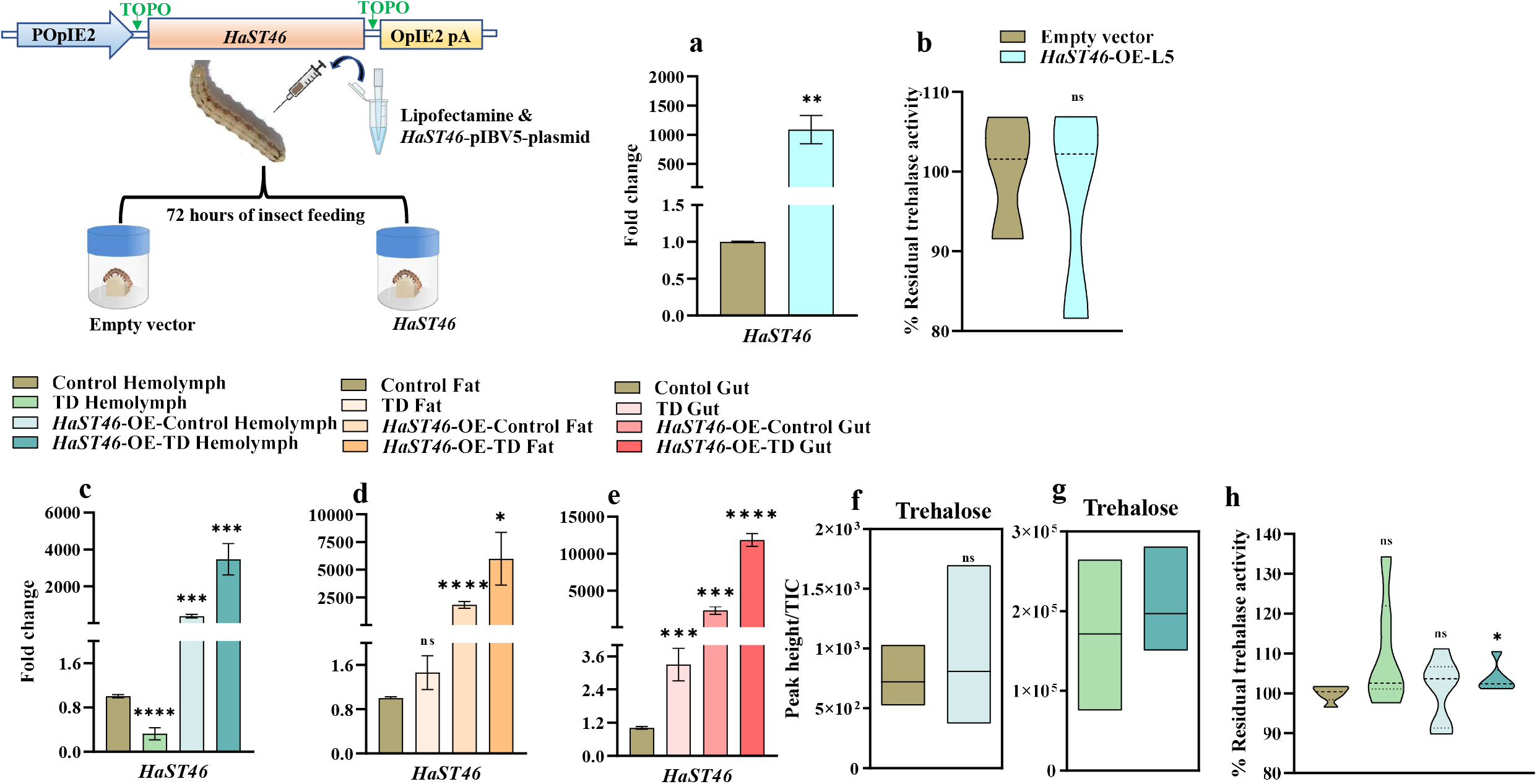
Overexpression of *HaST46* in fifth instar larvae of *H. armigera*. (a) qRT-PCR expression of *HaST46* in whole-body (b) In vivo trehalase activity in whole-body samples following *HaST46* overexpression. (c-e) Expression analysis of *HaST46* in haemolymph, fat body, and gut tissues under four feeding conditions: artificial diet, trehalose-rich diet (50mM), *HaST46* overexpression with artificial diet, and *HaST46* overexpression with trehalose-rich diet (50mM). Transcript levels were normalised to reference genes and expressed as fold change (2^^−ΔΔCt^). (f-g) LC–MS/MS-based quantification of trehalose in haemolymph after *HaST46* overexpression. (h) In vivo trehalase activity in in haemolymph, fat body, and gut tissues samples following *HaST46* overexpression. Data are shown as mean ± SEM, with significance assessed using an unpaired *t*-test (*P* < 0.05).

## 3. Discussion

Trehalose is key in energy homeostasis, functioning as a circulating sugar and substrate for rapid energy mobilization. The identification and functional characterization of *HaST46* as a trehalose-specific transporter localized to the posterior midgut, principal site for nutrient uptake, highlights its putative function in dietary sugar absorption **(Figure 1a and b)** (Terra et al. 2023). Furthermore, the nutritionally responsive expression patterns of *HaST46* suggest that its expression is coupled to the diet composition and larval metabolic demand. The higher expression of *HaST46* in complex or plant diet reflects an adaptive shift to facilitate the uptake of structurally and compositionally diverse carbohydrates in natural food sources **(Figure 1c and d)**. This adaptive plasticity aligns with earlier reports of the gut-specific modulation of nutrient transporters in response to dietary sugars in lepidopteran larvae (Azzouz et al. 2005), underscoring the ecological and physiological relevance of transporter regulation in herbivorous pests.

The substrate preference of HaST46 for trehalose over glucose represents a functional specialization **(Figure 1e and f)**. Despite structural conservation with GLUT1, HaST46 exhibits trehalose preference over glucose, suggesting an evolutionary divergence toward the hemolymph sugar composition of holometabolous insects **(Figure 1g)**. This specificity likely enhances transport efficiency and minimizes competitive inhibition, optimizing trehalose handling in sugar-rich dietary contexts.

At the metabolic level, the changes in trehalose biosynthesis and degradation pathways in response to transporter activity reflects a coordinated homeostatic system. The downregulation of endogenous trehalose synthesis in the presence of abundant dietary trehalose suggests a resource-efficient switch that curtails ATP-intensive biosynthesis in favour of direct absorption **(Figure 2a to e)**. Increased level of ATP compared to low level of ADP/AMP suggest low demand of ATP hydrolysis in exogenous supply of trehalose **(Figure 2f)**. This trade-off supports the notion that insects prioritize the least metabolically costly route to meet carbohydrate requirements, a trait advantageous for survival in fluctuating nutritional environments. Furthermore, systemic metabolic shifts, evidenced by increased glycolytic intermediates in trehalose-fed larvae, indicate that transporter-mediated uptake directly fuels energy metabolism, facilitating rapid developmental transitions. This catabolic reprogramming toward glucose-generating pathways mirrors the metabolic flux observed during feeding stages in other insects and provides a mechanistic explanation for the physiological benefits of dietary trehalose.

RNAi-mediated silencing reduces *HaST46* transcript abundance and triggers compensatory responses at both transcriptomic and enzymatic levels **(Figure 4a and b)**. Trehalose biosynthetic genes upregulation on *HaST46* knockdown implies compensation for decreased absorption by enhancing endogenous production. However, the simultaneous downregulation of other transporter isoforms suggests a response to reduced overall trehalose levels. Notably, despite transporter knockdown, nutritional indices remained relatively unaffected, indicating that *H. armigera* maintains systemic metabolic resilience, likely through alternative sugar transporters or compensatory physiological mechanisms. Conversely, *HaST46* overexpression did not disrupt homeostasis, suggesting that while the transporter enhances trehalose uptake, the system possesses regulatory checkpoints that prevent metabolic overload **(Figure 5a and b)**.

Together, these findings highlight role of *HaST46* in trehalose acquisition and regulation. Its dynamic expression, functional specificity, and involvement in feedback-controlled metabolic shifts position it as a key mediator of dietary sugar assimilation. Given that trehalose metabolism is unique to insects and absent in vertebrates, *HaST46* offers a promising molecular target for the selective disruption of energy metabolism in lepidopteran pests.

## 4. Materials and Methods

### 4.1. Differential gene expression and proteomic signature analysis

Publicly available *H. armigera* gut transcriptomic and proteomic datasets (Ioannidis et al. 2022) were used for expression profiling of *HaSTs*. Out of 5 tissue specific *HaSTs*, only *HaST46* and *HaST6*9 were mapped to gut transcriptomics and proteomics. For transcriptomics, raw fragments per kilobase of transcript per million mapped reads (FPKM) values were extracted, log_2_(x+1) transformed, and row-wise Z-score normalized before hierarchical clustering using Euclidean distance and complete linkage in the pheatmap package (R v4.x). For proteomics, the absence or presence of HaSTs in the proteomics set was denoted in binary form. Heatmaps were generated using TBtools 1.09 (Chen et al. 2020) to visualise stage- and tissue-specific expression patterns. For bar plots, mean FPKM values ± SEM were calculated across biological replicates.

### 4.2 *HaST46* overexpression in Sf9 cells

For overexpression, *HaST46* was cloned into the pIBV5 vector and sequence-confirmed. Sf9 cells (4×10^6) were seeded in a T25 cell culture flask (Corning, NY, USA) with 1 mL serum-free SF900 II SFM media (Thermo Scientific, Waltham, MA, USA) and allowed to adhere by incubating for 30 minutes at 27 °C. For transfection, 1.5 µg of either the EGFP_pIBV5 control plasmid or the *HaST46-pIBV5* plasmid was diluted in 100 µL SF-900 II SFM (without antibiotics). Separately, 8µL Cellfectin II reagent (Gibco, Waltham, MA, USA) was diluted in 100 µL Sf-900 II SFM and incubated for 30 minutes. The DNA and lipid solutions were combined (total 200 µL) and incubated for an additional 30 min to allow complex formation. The transfection mix was added dropwise while gently rocking the flask for an even distribution. Flasks were incubated for 5-7 hours at 27°C. After the incubation, the media was removed, and fresh Sf-900 II SFM containing blasticidin (Gibco, Waltham, MA, USA) antibiotic was added and incubated for 48 hours. Overexpression was confirmed by qRT-PCR. For the sugar uptake assay, 5mM and 10mM concentrations of glucose and trehalose were added to the media, and the cells were incubated for varying time points (0, 5, and 10 minutes). Following incubation, the media and cells were collected for metabolite extraction and analysis. Sugar uptake was determined from changes in extracellular and intracellular sugar concentrations by metabolite profiling.

### 4.3 Structural modelling and docking

Three dimensional structure of HaST46 was predicted using Alphafold 3 server (https://alphafoldserver.com/). Structural aligment of best model was performed using PyMol (The PyMOL Molecular Graphics System, Version 3.0 Schrödinger, LLC.) using Human Glut1 (PDB ID: 4PYP) structure as reference. Binding site residues of Glut1 were used to predict the binding pocket of HaST46 and molecular docking of glucose and trehalose was performed using AutoDock Tool as described earlier (Barbole et al. 2024).

### 4.4 Quantitative Real-Time PCR (qRT-PCR) Analysis

Total RNA was extracted from 80–100 mg of insect cells/tissues using the TRIzol reagent (Invitrogen, Waltham, MA, USA). To eliminate DNA contamination, RNA was treated with RQ1 RNase-free DNase. First-strand cDNA was synthesised from 5 µg RNA using oligo-dT primers and the High-Capacity cDNA Reverse Transcription Kit (Applied BioSystem, Foster, CA, USA), following the manufacturer’s protocol. Gene expression was quantified using a 7500 Fast Real-Time PCR System (Applied Biosystems, Foster, CA, USA) with Takara TB Green Premix Ex Taq II. Primer list provided in supplementary information (S3). Expression dynamics, including transcript abundance (2-ΔCt) and fold change (2-ΔΔCt), were visualised using GraphPad software. The expression of five *HaSTs, trehalose-6-phosphate synthase* (*TPS*), *trehalose-6-phosphate phosphatase* (*TPP*), and trehalase was assessed in trehalose-fed tissues, with the gut as the reference for fold change in fifth instar larvae. *HaSTs* expression was further evaluated in haemolymph, fat body, and silencing assays, using control expression as the reference.

### 4.5 Metabolite Analysis Using LC-Orbitrap-MS

To extract metabolites, 80-100mg of finely crushed sample powder was mixed with 500 µL of 80% MS-grade methanol to ensure efficient metabolite extraction (J.T. Baker, Center Valley, PA, USA). Methods were carried out according to the procedures described in the supplementary method 4.

### 4.6 Insect rearing

Larval populations of *H. armigera* were obtained from the ICAR-National Bureau of Agricultural Insect Resources (NBAIR), Bengaluru, India, and maintained on a chickpea-based artificial diet (AD) for multiple generations. Insects were reared at 25 ± 1°C, 70 ± 10% relative humidity, and a 16:8 h light: dark photoperiod. Adult mating boxes, containing equal numbers of males and females, were provided with 10% (w/v) sucrose solution and 1% (w/v) vitamin E, with a muslin cloth for egg collection (Dawkar et al. 2016).

### 4.7 Exogenous trehalose feeding and tissue collection for the bioassay

For exogenous trehalose feeding in the *H. armigera* bioassay, freshly moulted second instar larvae were fed AD supplemented with 10mM, 50mM, or 100mM trehalose for 6 days to assess the impact on nutritional indices efficiency of conversion of ingested food (ECI), efficiency of conversion of digested food (ECD), and approximate digestibility (AD) and expression of sugar transporter genes (*HaSTs*). Three biological replicates (n = 10 larvae each) were used, and the samples were flash-frozen in liquid nitrogen and stored at −80°C. For tissue collection, fifth instar larvae were starved for 1-2 hours followed by washing with phosphate-buffered water and dissected under RNase-free conditions to collect haemolymph, gut tissues (foregut, midgut, hindgut), and fat body. The samples were flash-frozen and stored at −80°C.

### 4.8 In vitro Targeted Putative Sugar Transporter Gene *HaST46* Silencing via dsRNA Feeding

Silencing of *the HaST46* gene was performed using microbial-based dsRNA production. To generate dsRNA targeting *HaST46*, a 400-bp fragment was PCR amplified using Phusion DNA Polymerase (Thermo Fisher Scientific, Waltham, MA, USA) and a Proflex PCR machine (Thermo Fisher Scientific, Waltham, MA, USA) with primers (**Supplementary File S3)**. Detailed methods are provided in **Supplementary method 2**.

### 4.9 Transient overexpression of *HaST46*

For overexpression, *HaST46* was cloned into the pIB/V5 vector (Thermo Fisher Scientific, Waltham, MA, USA) and sequence-verified. Larvae were starved for 3 h before injection. Further process were carried out according to the procedures described in the supplementary method 3. A mixture of 1500 ng of each plasmid and lipofectamine (Invitrogen, Thermo Fisher Scientific Inc., Waltham, MA, USA) (1:1 v/v), incubated for 30 minutes, was injected into the haemocoel between the 5th and 7th abdominal segments. The EGFP pIB/V5 plasmid served as the control (Barbole et al. 2024).

### 4.10 Enzymatic activity

The α,α-trehalase activity was determined by measuring the amount of glucose released from the hydrolysis of α,α-trehalose using the dinitrosalicylic acid (DNSA) reagent (Sigma-Aldrich, Merck KGaA, St. Louis, MO, USA). By monitoring the release of inorganic phosphate (Pi) from the crude extract using the malachite-green reagent (Sigma-Aldrich, St. Louis, MO, USA) to assess the enzymatic activity of trehalose 6-phosphate phosphatase (TPP). The assay was conducted following the modified protocol of (Klutts et al. 2003). Methods were performed as described in Supplementary method 4b.

### 4.11 Statistical analysis

All experiments contain 3 or more biological replicates. Data were expressed as mean ± SE using GraphPad Prism v8.0 (GraphPad Software, San Diego, CA, USA). The unpaired Student’s t-test. Asterisks indicate significant changes compared to control (*P-value < 0.05; **P-value < 0.01; ***P-value < 0.001, ****P-value < 0.0001).

## Acknowledgments

Fellowship support from CSIR-UGC-NET to BYC and VJN is acknowledged. ANRF (GAP336726) partially supports this work.

## Contributions

BYC designed and conducted the experiments, performed data curation, formal analysis, validation, and visualisation, and wrote and edited the original draft. VJN performed only transfection in the Sf9 cell line, whereas cloning, qRT, LC-MS sample preparation, and data analysis were performed by BYC. VTB carried out LC-MS experiments, and RSJ performed the analysis. RSJ conceptualisation, experiment design, supervision, provision of resources, funding acquisition, investigation, visualisation, docking, and methodology. BYC, VTB, and RSJ contributed to writing and revising the manuscript and approved the final version.

## Data availability statement

Data are available in the figures of the article and its supplementary materials.

## Supplementary information

**Supplementary information S1**: H. armigera STs gene sequences

**Supplementary information S2:** Name of putative HaSTs and its corresponding gene id from NCBI gene database and scaffold id from transcriptome data

**Supplementary information S3:** HaST primers used for real-time PCR analysis and silencing experiment

**Supplementary information S4:** General methods for real-time PCR, TPP and tre-halase enzymatic acitivites.

**Supplementary information S5:** *H. armigera* growth and phenotype upon silencing and upregulation. Also real-time expression analysis of HaSTs related.

