## Supplementary material for "Trehalose Transport Dynamics Underpin a Metabolic Trade-Off between Exogenous Uptake and Endogenous Synthesis in Lepidopteran Insects": Supplementary File_S1-Gene sequence-Ha Only.docx

***HaST09***

**>evm.model.scaffold_139.12 gene=HaOG203721_tret09**

ATGAACAAAGGAAGACTTAGGCAGTGTCATGCAGCCATGGCTGGCGGCTTCGGGTCGCTGGTCATGGGCGCTCTCAACGTCTGGCCCTCCTACACTTCAGAGCTCTACGCGTCCAACACCACTACTCCCCTATCTGCGCCTATGACGAAAGGTGAAGAGGCTTTGTTAGGGAGCCTGCCTTCGCTAGGAGCGATTCTCGGCTCAGCTGTAGCTGGGACACTGATCAATATGTTCGGGAGACGAAATGGTGGTGTTATATTGTCTTTGCCTTGTGTGATGTCCTGGGCAATAATAGGGGTAGCGAGTTCTATGAACCTGATTCTGGCAGCTAGATTCTTGTCCGGTATCACGGGAGGCAGCTTTCTAGTACTGACGCCTATCTTCATATCAGAAGTGGCTGAAGACGCTATTCGCGGAGGCTTAGCTTCAGCATCAATATTCCTATACTGCACGGGCACTCTACTCTCGTACATCGTGGGCTGGTGCCTCACGTACAGGACTATCATCTGGCTTCACTTGTTACTCAGCGTGTTGTGTACTCTACTTATACTGCTGGTGGTTGAAAGTCCCGTGTATCTGCTGCGGCAGAAGAGGGAAGAGGACGCAAGAGAAGCGATAGCGAAATACAGAGGAGTTCCAGTATCGTCAATGGTTGTGCTCGATGAACTGACGCGGATGAAGCAACAGATCATGCCGGCTGTCGAACTTGTGTCTATTACAGATATGGATCCCAAAGCAGAAGAAGCTGAAAAGCAGAAACTAAATAATGAAGAAGAGATCGTCCAAGAACAGCCTAAATCGATGTCACCGATCAAGTTACTGTTCGTATCACCGGTATCGCGGCAGGCGTTCATAGTTGTAATGACAGTCATCACGATGCAGGTATTCATGGGCATAGTTCCCGTACAAGTGTACGCTAAGACAGTGTTCACGGAGACAGACCCCAGCAAGTCAGACCTCTATACCGTCGTGTTTGCTGTCATACAGTTCTTTGGAGCTTTGACCTCTTCGCTTGTAGCTGATAAGGCTGGTAGACGGATTCTCATCATCATCTCGTCAATCCTCGTATTCCTCTGCATGGTGTCCCTCGGCTTCCTCCTGCAGACGCGCATGGCGCCGGCCTGGCTGACGGTGGTGCTGCTGATGCTGTACTGCTTCTCATTCCTCATAGGAGCTGGAAGTATACCCTATGTGTTGTTGGCTGAGGTCTTTGAGTCACAGGTCCAAGGCCTAGGTTCAATGATCGTCATAGAATGGGTTTGGTTCCTCAACTTCTTTATTCTAGGTATCTTCCCATATATGTTGAGTGCTCTCAAAATCCACGGGGCCTTCTATTTCTTTGGTGCCATGAGCCTTCTAAACGCAATATTAGCGTTTATACTAGTGCCGGAAACTAAGGGACTGTCAAATGCACAAATTCAGGATCTTTTTAGATCAAGAAAGAAGAATTAG

***HaST29***

**>evm.model.scaffold_227.8 gene=HaOG207130_tret29**

ATGGATTCGTATTTGAAGACACAGATTTTAATAGTGGCCTGTGTAAACATAGGCCAGTTCATAGACGGCTACAGTGTTGGCTGGTCAGCTCCCATCATCCCCAAGTTACAGGATCCTCAGCAGTCACCCCTGCCTGAGGTAGTGACTGACTTCCAGGTGGCCTGTATAGGATCCTTGCTGTATATTGGAGCTATAGTTGGTCCATATATCCCAAGCTACCTATCCAATGTAATAGGTCGGAAACCGTGCCTCCTACTAGGCGGGCTACTGAACCTTCTCGCCATCATCCTCATCATCACCACTCGTAACATCGCCATGGTGTATGCTGTTAGAATCATCAGCGGATTGGGGATGGGGATGGTGACAGTCAGTAACTTGGTGTACATTGGGGAGATTGCGTCTACAAACATCCGTGGCATACTCCTAACATCGACATCTATTGTTGGCATATTCGGGACGCTAGCAGCCTACAGTATAGGACCTTATGTGTCCTATGAAGCAACTGGTTATATAGCTCTGGTTATTAATATAGTTCATGTGATCGGTATATTGTTTATACCGGAGTCTCCTGTGTATTATGCTTTGAAAGGTAAAGAAACAGAAACCAAATCAACACTCCGACTTCTCGGCCGACTGGATGACTTAGAAAACGTCTTCGAGTCAGTTCAAGACTTGGACCCTAATGATGGCCACAGTTGGAAGTCATGGGTGAAGATATTCACTGTCAAGTCCAATAGAAGGTCCCTGATCATCACGTTTAGTCTGCTGACTCTTCAGCAGATGAGCGGAGTGGCAGCTGTGCTGTTTTTTGTGACTACCATATTTCAGCTGGCTGGGTCTTCGATACGGCCAGATCTAGCCACAATCCTGGTAGGTGCAACACGTCTTCTATCTAGTTTGATAGCACCAACGCTGGTGGAGAGAGCTGGCAGAAGAATCTTGCTTCTAACTTCCACTGCATTTTGTGCTATCAGCTTGTTTATCCTCGGCACATATTTCTATTTGGATCGGATACAGAGTTCCATCATTTCGGACATCAGATGGCTGCCGCTCATGGCTCTAATTATGTACTTCTTCTCTTATGAAAGTGGTTTCGGCACAATGCCCTTCGCCCTCGTCGGCGAGATGTTCAAGGGCAACGCCAGAAGTCCTGGCTCCGCCATCTCCATGACCACCGCTTGGCTCATCGGCTTCCTCATCGCCACCAGCTTCAACACCATGCTGAACAGCATCGGCAGTGACGTCACGTTCCTCGTATTCTCTCTCTCCTGTGTCCTGGCTTGCTTGTTCACTTACAAGTTTGTTCCTGAGACTAAGGGGAGGACTTTGAGTGAGATTCAGCAGATTTTGAGTGGGTGA

***HaST46***

**>evm.model.scaffold_34.66 gene=HaOG210281_Tret46**

ATGTGTGATGCAGTGTCACGGAAACATTCTGAGATGGGTGACGGAACAGTGCCAGAGTTAGAGAAAATCCATAAAAATGGAACAACGGACAATGGGAAAGAACTTAGTGCTTCTAATGATGAATACCTCAATAACCAACGATCCCCGTTTAGGAGACAGGCTATCATCTCATTTGGTGTGTTCATGCTCACCCTCGGGGCTGGTGCCACGTCAGGCATCTCTGCCATCTTAATTCCACAACTACAACATGCAAAGGGGAAAAAAGCCTTCTCAGTAGAATTGGTATCATGGGTAGCTGCCATATCATCGTTGGCTCTCTTCTTCGGTAACCTGATGTCGGGATATTTGATGGATAGACTTGGAAGAAGAATGTCTAGTCTACTTCTGGCGGGCACATTTGTGGCTGGCTGGTTGATTATTGGGTTCTCAAATGACCTCATGTTCCTAATCTTAATAGGAAGATTTATCACGGGGTTATGTCAAGGGTGGCTTGGCCCCCTTGGCCCAGTCTACGTAGGAGAATTTAGTAGTCCTGCTTACAGAGGACTGTTCTTAGCAGCTTTATCTTTAGGAATAGCAGTGGGTGTTTTCATGTCACATTTATTTGGCACTTTCCTGCATTGGAGTATATCGTCCTTGCTTTGTGGATTATTCCCACTGATTGGTGGTGTTATACTTTATTACGCACCAGAATCACCTTCGTGGTTAGCGTCTAAACAGCGAATTGATGAATGCATAATCTCGTACCAATGGTACAGAGGAAACAGTGCAGCTATGAAAACTGAACTTGATAAAATGATTGCTGATATTTCAGCGAAAGGCAATAATCAGAGTAAATTGCAAATTATAGCAGCGAACATAAAGAAGCCAGAGTTTTATAAACCATTAGGTATTATGACGACCTTCTTCGTTATAACGCAACTTTCTGGTGTCAACGTTATTTGTGCGTACACTACAGAGATGATGAAGGAACTCATTGGTAGTGGTTCGACTAGTTCACACGCCTACGCCGCTATGTTAAGTATAGACGTGTTGCGATGCGTGTCACTCGGCGCTGCTTGCATTATGCTCAGGAGATCAGGTAGAAGGCCCATGGCTATATTCAGTGGAGTATTCACATCGTTATCTTTAATTTCACTAGCTTTGTACCTGTATCTCAACGACTCTGGTGTCATCCATCACATATCACCATTCATTTCATTAGGTTTAATGGCATTCTACATAGTAGTATCCAATTTGGGAATATGTCCACTGCCATGGAATATGGTTGGTGAACTATTTGCCGTCGAAACTAAAGGAGTGTGTTCAGGCATTAGTGTCATGATGACTTCTGTGGCTTTCTTCGGAGTAGTGAAGACGGCGCCGTCCATGTTTAGAAGTATAGGTCACCATGGAACGTATCTCTTCTACGGGTTATCCACCCTGTGTGGTACCATATTCTTATACTTATGTTTACCCGAGACAAAAGACAAAACTTTGTTACAAATAGAAGAACACTTTAGGTATGGCAAGAAAAAGAATGATAGCAAAGAAACAGATAATATTTAA

***HaST46*_ds-sequence**

TGAGATGGGTGACGGAACAGTGCCAGAGTTAGAGAAAATCCATAAAAATGGAACAACGGACAATGGGAAAGAACTTAGTGCTTCTAATGATGAATACCTCAATAACCAACGATCCCCGTTTAGGAGACAGGCTATCATCTCATTTGGTGTGTTCATGCTCACCCTCGGGGCTGGTGCCACGTCAGGCATCTCTGCCATCTTAATTCCACAACTACAACATGCAAAGGGGAAAAAAGCCTTCTCAGTAGAATTGGTATCATGGGTAGCTGCCATATCATCGTTGGCTCTCTTCTTCGGTAACCTGATGTCGGGATATTTGATGGATAGACTTGGAAGAAGAATGTCTAGTCTACTTCTGGCGGGCACATTTGTGGCTGGCTGGTTGATTATTGGGTTCTCA

***HaST64***

**>evm.model.scaffold_70.36 gene=HaOG215283_tret64**

ATGAGTTTCAATAAAAACAACCCCAACGCCATGGGGAAGATCATGGGCTACATCAAGCAGCTCTCCACCGAAGTGGGTGGGAGCGAGCAAACTCGCCGAGGTCAAGGCGACGAGGAGAGGTTGTACCGGACCCGAGGGCCCAAGTACTCTAGAGTCCCATCAAGACCGACGCTTTCTGCTTCTACGACCTGTACCTCTCTGGCAGAATCTTGTGGGTCCCAAGGGACATTGGTGCCTAACTACGCAACAATCCCAGAAACAGTTTCCACTGAAAGCAGCAGTGAAGACGAGCAGGACTCATTCGAGAACACCCGTCGCCATTTCCAACAACTTCGGCAGATCAGTCTAGGAAACGAGTTTAAGTACAAGATGGAGATGGAAATAAAGAGTGCGAAGGAGGAGAATTTGAGAAATTCGATTCCTTTTGTCAAACAACTGAGCACTGACAGCAGTAAAGTAAAACCGGACTATGCAATCAACGGGGATACTCCACCATATGCTCCAACAACCCAACGACTATTTCTGTGGACACAACTTTTGGCCGCATTCGCTGTGTCTATGGGTTCGTTGATTGTCGGCTTCTCGTCCGGCTATACTTCTCCTGCATTGATAAGCATGAACTCTACGCTTCACATGACTAAAGAAGAGTCAACATGGGTCGGCGGTCTTATGCCTCTGGCTGCATTGGTTGGTGGAGTCGCAGGAGGACCTCTGATAGAATGCATTGGAAGACGATGGACCATAATGGGAATGGCTTTACCATTCTTCCTCGGCTGGATGCTCATAGCAACTGCGTCAAACGTGCTAATGGTGTTCGCTGGAAGAGTTTTCTGCGGAGTGTGTGTCGGAATAGTCTCCCTGGCATTCCCAGTTTACCTCGGTGAAACGCTACAGCCCGAAGTACGAGGTGCGTTTGGATTGTTGCCTACTGCCTTTGGTAACACTGGAATACTTTTATCATTCTTTGTGGGAAGCTACCTTGACTGGTCGAAACTAGCATTCTTTGGAGCTGCATTACCGGTACCATTCTTCCTGCTCATGCTGCTTACGCCTGAAACCCCACGCTGGTTTGTGTCCAAAGGACGCCCTGAAGATGCTCGTAAAGCGCTTCAATGGCTTCGAGGAAAAAATACAAACGTTGACAAGGAAATGAAGGATCTTACACGTACACAGGCTGATTCGGATAGAACAGGAGGAAATGCTTTCAGACAACTTTTTACTCTTAAATACATGCCCGCTGTCCTTATTTCTCTTGGATTAATGTTGTTCCAACAGTTAAGTGGTATTAATGCAGTAATTTTCTACGCCGCGTCAATCTTCAAAATGTCTGGAAGCACTGTTGACGAAAACTTATCTAGTATCATAATTGGAATCGTCAACTTTGTTTCTACATTTATTGCCACAGCTATCATTGACCGCTTGGGACGTAAAATGTTGTTATACATTTCCTCAGTTTCTATGATAGTTACTCTAGTTTCACTGGGAGCTTACTTTTATGTGATGGATTCAGGAGTTGATGTCACGGCTTTTGGATGGTTGCCACTTGCTTGTCTTGTCATTTATGTATTGGGATTCTCTATTGGCTTTGGACCCATCCCGTGGCTCATGTTAGGTGAAATTCTACCATCCAAAATCCGTGGCACAGCTGCATCTCTTGCGACTGGATTCAACTGGACGTGTACCTTCATTGTCACTAAAACTTTCCACAATATCATCGACGCCATTCATATGTACGGTACAGTGTGGCTGTTTGCTGTCATTTGTTTAATTGGGCTGTTTTTCGTAATATTCTTTGTCCCTGAGACTCGAGGTAAAAGTTTAGAGGAGATTGAAAGGAAATTAACAGGGGGTTCACGAAGAGTGCGGCATATTAGCAGCAGTAAGCAACCACAAAATGGCTGTTAA

***HaST69***

**>evm.model.scaffold_8.129 gene=HaOG215993_tret69**

ATGAGGCTCACGCGGCGCAGGTGGGCAGAACTCCAAATATTCCGAACAAAAGCTACGCTTATCACCGCTACAGCGGGTACCTGCTACGGGTGGCCCTCACCTACTCTACCGTACCTACTATCCGAAGAGAGTTCAATCAAAACAACAGCTGACGAGGGATCATGGATAGTCTCGATAATGATCCTGTGCTCCGCGTTGACGCCTGTTCCCTCCGCCTACTTCGCCGACCGGTTCGGCAGGAAGACCACGCTACTCCTCGGTGCGGTGCCGTTCATCCTGGGTTGGGTGCTGGTCATCGTGGCCAACTCCGTCGCTCTGCTCTACGTGGCGCGGATGTTCTCCGGCTTGGGCTATGGAATTGTCTACACAGTTGCTCCAATGTACACAGGAGAAATCGCTACCAATCAAGTTCGAGGAGCCCTCTCCACACTCATCACGTTAATGAATAAAGTCGGAATTCTTGCCCAGTACTGCATCGGTCCGTTCGTCTCGATGCAAACCCTCGCTGCCATCAACTTGATCCTGCCCGTCACATTTGTCATCACCTTCATCTTTTTGCCAGAATCTCCTTACTACTACTTAAAATTTGAGCGAAGTGAGAGAGCCGAGCGCTCACTCAGGAATCTACGCAGTGGCGACATTAGAACTGAACTCAAAAGTATAGAACTGAACGTTCAAGAAGACATGAAGAATAGAGGAACTTGGGGAGACTTGATCACTGAAGCTACCAACAGGAAAGCAATGTGGATTACGCTTGGTATATTCACGATACAGCAGCTATGTGGCAGTGCTGCTGTGGTGGCATACGCGCAGGTCATATTCAACTGCACGACCAGCCCAGTCACTCCAAACATTACTGGAGCAGAAAATGTTACCGCTTCCGCTTCTATTGAACCCTACCAAGAATCTATTATTCTTGGTTGTGTACAAGTAGCCACCTGCGTCCTGTCAGTAATACTCGTCGACCGTGTTGGTAGAAAGCCCCTTCTGTTGCTATCGGCTCTTGGAGTGGGCCTTATGAACGGCACAATTGGAACATACTTCTACTTCGACCATGTCAACAAAGAAGCCGTTGCACATCTTCACTGGATACCCCTCGCCGCTCTTCTCGTTTACATCGTTTGCTACGCCATCGGGTTGTCGACGGTACCCTACGTCATCATAGGAGAGATGTTCCCGACCAACGTCAAGTTGTACGCTTCCTGTATCGCTCACATCTACACCGGCGTCTCCATGTTCGCTGTTCAAAAACTATTCCAGGTGGTCAAAGACGCATATCAAATCTACACAGTATTCTGGGGATTCGCCACGTTCTCGCTGCTGGGGCTGGTGTTCATGCTGATCATGTTGCCGGAGACGAAGGGCAAGTCGTTCGCGAGCATCCAGGCGCAGCTCAAGCGGGAGGTGGCCCGCGACAACGCTAAGAAACTGGCCACCGTCGAATACTGA

**EU878265.1 (159-2639_cds) *Helicoverpa armigera* trehalose 6-phosphate synthase isoform I mRNA, complete cds, alternatively spliced**

**>mRNA_tps_aa(6-507)final_nt(174-1679)**

AGCAGTGCCAGTCGATCCGCGTGCAACAGCAAGGGAAGCATGATCGTTGTGTCGAACAGGTTGCCCTTCATCCTCAAGAGAAATGACAAGACTGGCGGTCTGGAGAGGAAAGCCAGTGCTGGTGGGTTGGTGACAGCAGTAGCTCCAGTGGTGATCCGTGGAGGCGGCATCTGGGTGGGATGGCCAGGCATACATCTGGATGACCCCAATGAAAAGATCCCGGAGTCAGACCCCAACGACAAGACCCCTACTGCCGGCTTGCTATCAGAGAAGATAGTCCCCGTGCACGCTGAGCCCAAACTCTTCGACAGCTACTACAATGGCTGCTGCAACGGTACTTTCTGGCCCCTCTTCCACTCCATGCCTGACCGAGCCACCTTCATCGCTGACCACTGGAGGGCATACATCAAGTGCAACGAGGAGTTCGCTGAGAAGACAGTGTATGCTCTGCATCTGCTCAAACAACAGAAGGGGAAGAATGGGACCTCTCCACCGATCGTGTGGGTCCATGATTACCATCTTATGTTAGCTGCTAACTGGATTAGACAGCGAGTTGAGGAAGATGACATAAAATGCAAGCTTGCGTTCTTTCTGCACATTCCTTTCCCCCCGTGGGACATATTCAGGCTGTTCCCATGGTCTGATGAAGTATTGCAGGGCATTCTTGGTTGTGACATGGTCGGATTCCACATAACTGACTACTGCCTGAACTTCATTGATTGTTGCCAAAGAAACTTAGGTTGTCGTGTGGACAGAAAGAATCTGCTAGTTGAATTGGGAGGTCGCACCATCTGTGTCCGACCGTTACCTATCGGAGTACCCTTCGACAGATTTGTCCAGTTGGCTCAAAACGCAAAGACAGTGCTCTCTACAAGCCAACAAATTATATTAGGAGTTGATAGACTGGATTATACCAAAGGATTAGTACATAGACTGAAAGCTTTCGAGAGATTACTGGAAAAGTATCCCGAGCACATCAAGAAAGTAATGCTGCTTCAGATCTCGGTGCCGTCAAGAACGGACGTCAAGGAATACCAGGACTTAAAGGAAGAGATGGATCAGCTGGTTGGAAGAATAAACGGAAGATTTACTACTCCAAACTGGTCACCTATTAGGTACATTTACGGATGCGTCGGCCAGGATGAACTAGCTGCCTTCTACCGCGATGCTGCAGTAGCCCTGGTTACACCTCTGCGAGATGGCATGAACCTCGTCGCTAAGGAGTTCGTAGCCTGTCAGATTAACAAGCCTCCAGGAGTGCTGATCGTGTCACCCTTCGCCGGTGCTGGAGAAATGATGCACGAAGCTCTCATCTGTAATCCGTATGAATTGGACGATGCTGCTGAAGTCATTCACAGGGCGCTGATAATGCCGGAAGATGAGCGCACAGTCCGTATGAACCACTTGAGAAGACGTGAGCAGCTCAATGATGTTGATAGCTGGATGAAGGCGTTCTTGAAAGCCATGGAC

TCTTTGGAAGAGGAGGCTGATGATATTGGTGCCACG

**>mRNA_tpp_aa(512-783)nt(1692-2507)final_nt(1681-2529)**

CCATGCAGCCTGTCACCATTGATGACTTCGATGAATATCTTTCTAAGTACATTGGCTACACACAAAAGCTGGCATTACTACTTGACTACGATGGTACTCTAGCCCCCATCGCGCCTCACCCTGACCTGGCAACCTTACCCTTGGAGACCAAGCATACTCTGCAGGGGCTGTCCAATATGTCCGATGTCTACATCGCCATCATCTCCGGCAGAAATGTCAACAACGTTAAGAATATGGTTGGCATTGAAGGCATCACGTACGCTGGTAACCATGGTCTGGAAATCCTGCACCCAGACGGCAACAAGTTCGTTCATCCCATGCCCATGGAGTTGCAGGACAAAGTCGTCGACCTGCTCAAGGCTTTGCAAGAACAGGTGTGCAAAGACGGAGCCTGGGTAGAGAACAAGGGAGCTCTCCTGACGTTCCACTACCGCGAGACGCCGGCTGACAAGCGGCCGGCGCTGGTGGAGCAAGCCCGCAAGCTGATCACGGCGGCTGGCTTCACGCCCGCGCCTGCCCACTGCGCCCTCGAGGCCAGGCCGCCCGTCGAGTGGGATAAGGGTCGTGCATCCATTTACATCTTGAGGACAGCGTTCGGTTTGGACTGGAGCGAAAGGATTAGGATTATCTATGCTGGTGATGACGTCACCGATGAAGACGCCATGTTGGCCCTCAAAGGTATGGCAGCTACATTCCGCATCGCTTCATCCCAAATCACGAAGACATCAGCTGAACGTCGTCTATCCTCCACGGGCTCAGTACTGGCCATGCTCAAATGGGTGGAACGTCACTTTTCCCGCCGCAAGCCGCGCGCCAACTCGTTGACGTACAAAAGCGCGCGAAAGGCCA

***Ha-trehalase-1***

**>ENA|KJ652557|KJ652557.1 *Helicoverpa armigera* soluble trehalase mRNA, complete cds.**

GCAGTCGTATCCGCGATCGCGACTCGTGCGCATGTAATCGATTATTCCGACGAAATGTCGAATCGATAAAACAATCGATAAACATTGATACGATCGTTTAGTTGTACCAGTGAAATAGAGTAATATTAGTTTTTGTTTATTAGTTTATTCGAGATTTTATCGATATATCGAGTTGTCTGGACGAGTTGTGATTTGTAAGCTGTGAAGTGAATGGTGTCCTTTTGTAAGATGCGAGAACTCCTGATCTTGTTGGCGGCCGCGGCTGGGCTGGCCAGCGCTGACCTGCCACTCACCTGCACCAAACCCGTCTACTGCAACAGCAACCTGCTCCATCAAATCCAAATGGCGAGGCTCTACAATGACTCCAAGACCTTCGTAGACCTTCAAATGAACTTCGATGAAAACAAAACTTTGACCGACTTCGAAACCTTTTTCAACCTTCATAACAAAAACCCGACTAAGGAACAGTTGATGGAATTCGTCAATGAATACTTTTCCAACGACAACGAACTGGAGCCATGGCAGCCAAAAGACTTCAGTGACAATCCAGCATTTCTTGCTAAAATAAAGGACGATGCGTTAAGGGAGTTTGGAAAAGGTATCAATAACATTTGGCCACTTTTGGCACGGAAAGTTAAAGCAGAGGTGTTCCAGAAGCCCGATCAATTTAGTTTAGTACCCCTGACTCATGGATTCATAATACCCGGTGGACGATTCAAGGAAATCTATTACTGGGACACTTTCTGGATCATTGAAGGTCTTTTGATAAGTGGAATGCAA

GAAACCGCTAAAGGAATGATTGAAAATCTCATTGAATTATTGAATTTATTTGGCCACATCCCTAATGGTAGCAGAGGGTATTACCAGCAACGTAGTCAACCTCCTATGTTAAATGCCATGGTGGCTACTTACTACATGTATACCAAAGATCTCGAATTCCTCAGAAATAACATCGCATACTTAGAAAAAGAATTGGACTTCTGGATGGATAATAGAGTGGTATCAGTTAACAGAGGAGGTAAAAATTATACGCTTCTTAGATACTATGCCCCAAGCAAAGGCCCTAGACCCGAATCGTATTATGAGGACTACAGCAACACTGAAGGTTTTTCGGAAGAAGACAGTACCAATTTCTGCATCGATATCAAAAGTGCGGCTGAGAGCGGGTGGGACTTCTCAACGCGTTGGTTCCTCATGCCAGACGGCAGTAACAATGGCACTTTAACTGATCTGCACACGCGGTACATCATACCCGTTGACTTAAACGCCATCTTCGCCGGAGCTGCCCAGTACGTGTCAAACTTCCACGCCCTCTTAAAGAACCCGCAAAAAGCTGCTAGGTACGGACAGCTAGCACAAACCTGGAGAGACAACATTCAGGCAGTGCTGTGGAACGATCAAGATGCGATGTGGTACGACTTCAATATTAGGGACAATTTACATCGCAGATACTACTACTCGTCTAACGCTGCGCCGCTATGGCAGAATGCCGTTAATCCAGATTTTCTGAAACTCAATGCTGACAGGATTTTGAAAGCTATCACTGAATCCGGAGGTGTAGACTTCCCCGGAGGTGTACCCACGTCGCTCATCAGGAGTGGAGAGCAGTGGGACTTCCCCAATGTGTGGCCTCCAGAGGTGAGCATCGAAGTCGCTGCTATTGAGAATATCGGGACGCCTGAGGCTATTACTTTGGCGCAGGAAGTAGCACAAACTTTCGTGAGGTCTTGTCACTGGGGCTTCCAGAAGTACAAGCAGATGTTTGAGAAGTACGATGCCGAGACGCCCGGCAGGTTCGGCGGTGGCGGTGAATATAATGTGCAGTTCGGTTTTGGTTGGAGTAACGGCGTCGTACTGGAATTTCTAAATAAATATGGGTCTCAGCTAACAGCCGACGACTCTAACAATACGAATAATAGTGCATGACTAGGCGTGGATATTGTGAACAATTTCCTAATTTTCATATCCACTATTAAAATACTTATAGCTTCCTACATTAGTACATAATATTTGAAGGTTGTGAACTTACGATTTTGACTAAATTAAGTGATTCTGACTAAAAACGGTGCCATCACTAATGAAATTTATATAAAAAAAAAATAAAACCGTGTGATGTCGGATAATGATAGAACATTTTACTCTACAACAGCGTTATACAGACAGGCTGTAAGAAACTCAGGTGCTTCAGACCAGGCAGAATATATTTCCCAATACTTAATTTAATAGAACAAAAAAAAAAAAAAAAAAAAAA

***>Ha-trehalase-2***

ATGGATCGGAGTCACTTGCCACCGACCTGTTCTAGCACCATCTATTGCCACGGGCCCCTACTAGACACGGTACAAATGGCGGGCCTGTACAACGACTCTAAGACCTTCGTGGATATGAAGCTCAAGCTGTCTGCCAACATCACCATGGAACACTTTCAGGAGATGATGGCCAGGACAGGTTCACACCCGACCAAGGCTGACATCCAGGAGTTTGTCAATCAGAACTTCGACCCTGGGGGCTCCGAGTTCGAAGACTGGCGGCCTACTGACTGGAAGGATAATCCTGCATTTCTGCAAAACATCAAGGATCCTCTGCTCCACCAATGGGCTGCAGAGCTGAACAGACTGTGGTTACAGCTTGGCAGGAAGATGAAGCCGCATGTGAAGAACAACCAGGATCTGTACTCTATTATCTACGTGGATAATCCGGTTATTG

TGCCTGGTGGTCGTTTCCGAGAGTTCTACTACTGGGACTCCTACTGGATCATCAAGGGTCTGCTTCTGTCCGAGATGAGGGCCACAGCTAAAGGCATGGTGTCAAACTTCATGGATATTGTGGAGAGGATCGGCTTCATTCCCAATGGGGGGAGGATATATTATGCTATGAGATCACAGCCCCCACTCCTAATCCCCATGGTGAAGATAATACTGGATGATATGGACGACCTGGAGTACTTGCGCCAACACATACACACCTTAGACAGAGAGTATGACTACTGGATGACTAACCATACTGTCGAGGTGGACCATAATGGGCATAGATACACGCTAGCGAGGTATTACGATCAGTCACAAGGACCCAGGCCTGAGAGTTACAAGGAAGACGTCGATGTGGCTAGACACTTTGACACAAATGACAAGAAAGAGGAGTTATACGCCGAGCTGAAGGCGGCTGCTGAGCCAGGATGGGACTTCTCATCCAGGTG

GTTCATACTCAATGGCACCAATAGAGGTAACCTAACAAACCTGAAGACCCGCTCCATTATCCCGGTGGACCTCAACGCCATCATGTGCTGGAACGCACAACTCCTGAGAGACTTCCACCTCAGGCTCGGCAATATCGATAAGGCGGAGTACTATAGGAACGTTCATGCGAGGTTCATGGATGCTATTGAACAGGTCCTATGGCACGAAGACGTAGGAGTCTGGCTAGACTACAGCCTGGAGTCGGGCAGACGCCGCGATTACTTCTACCCGTCAAACGTGTCGCCTCTATGGGCAGTCTGCTACGATCAGGCCAGAAAGGACTACTATGTCAACAGAGTTGTTAACTATCTGGATAAAGTTAAAGTGGACATTTTCGACGGCGGCATCCCAACAACTTTCGAACATTCTGGAGAGCAGTGGGACTACCCGAATGCCTGGCCGCCATTACAGTACATAGTGGTAATGGGCTTAGCTAATACTGGCCAGCCA

GAGGCTGTGAGACTGGCCAGCGAGATCGCTACGAAGTGGGTGCGTTCGAATTTCGAAGTTTGGAAACAGAAGACTGCTATGCTTGAAAAGTACGACGCGACAATTTTCGGCGGTCTCGGCGGAGGCGGCGAGTACGTTGTACAAACAGGCTTTGGTTGGACCAATGGCGTGATCATGGCCATGCTCAACAAATGGGGAGATACGCTTACTTCAGCGGACGCGTTCGGGACGGGCGTGACGGCTGACTCCGGTGCTGTGTACGGAGCGCATGTCGGCGCTAGCGGCGTGGCCACGGCTATTCTAGTAGTGCTCGCATCTTTGGCTGCGGGGACCCTTGGACTCATCGTATACCGAAAACGCAGGGACTACATCCGAGTATCAGGGGGCGAAGACTACAAATTGCTCTCCCGAAGACCTTAC
