## Supplementary material for "Trehalose Transport Dynamics Underpin a Metabolic Trade-Off between Exogenous Uptake and Endogenous Synthesis in Lepidopteran Insects": Supplementary File_S4_Genetics_1192025.docx

**Supplementary methods**

**Supplementary method 1**: Quantitative Reverse Transcription Polymerase Chain Reaction (qRT-PCR)

The reaction cycle used for real-time PCR was as follows

| Temperature | 95°C | 95°C | 60°C | 95°C | 60°C | 95°C |
| --- | --- | --- | --- | --- | --- | --- |
| Stage | Stage 1 | Stage 2 | | Stage 3 (Dissociation stage) | | |
| Time | 30 sec | 5 secs | 45sec | 15 sec | 15 sec | 15 sec |

**Supplementary method 2**

**In vitro Targeted Putative Sugar Transporter Gene *HaST46* Silencing via dsRNA Feeding**

PCR conditions included 98°C for 30 s, followed by 35 cycles of 98°C for 10 s, 60°C for 30 s, and 72°C for 30 s, with a final extension at 72°C for 10 min. The purified PCR product was cloned into the pGEM-T Easy vector (Promega Corporation, Madison, WI, USA), excised, and ligated into the L4440 vector using NotI and EcoRI. The ds*HaST46*-L4440 plasmid was transformed into E. coli HT115 competent cells. A single colony was cultured in a 5 mL primary culture with tetracycline and ampicillin. A 200 mL 2x YT media with antibiotics was inoculated for secondary culture and induced with 1mM isopropyl β-D-1-thiogalactopyranoside (IPTG) at an OD of 0.4. After 4–5 h at 37°C, the cells were harvested, resuspended in nuclease-free water (OD 2.0), and mixed into the larval feed. A bioassay with 15-s instar larvae per treatment was conducted for 6 days using an empty L4440 vector as a control. The feed was replaced, and the parameters were recorded every alternate day. On day 6, seven larvae per treatment were starved for 2-3 h, flash-frozen, and stored at -80°C. Nutritional indices (ECI, ECD, ADI) and body weight changes were calculated ^37^. RNA was extracted using TRIzol, treated with RQ1 RNase-free DNase, and visualised on a 1.2% agarose gel to confirm ds*HaST46* production. qRT-PCR was performed to assess *HaST46* expression in the control and treated insects, using actin as a reference.

**Supplementary method 3**

**Transient overexpression of *HaST46***

A mixture of 1500 ng of each plasmid and lipofectamine (Invitrogen, Thermo Fisher Scientific Inc., Waltham, MA, USA) (1:1 v/v), incubated for 30 minutes, was injected into the haemocoel between the 5th and 7th abdominal segments. The EGFP pIB/V5 plasmid served as the control. Gene expression was quantified by qRT-PCR at 48 and 72 h post-injection, with six biological replicates (three larvae per replicate). For tissue-specific expression analysis, the tissues were dissected 48 h after injection. Following injection, larvae were maintained on AD supplemented with 50mM trehalose- or control AD for 24 h, then flash-frozen, and stored at -80°C

**Supplementary method 4: Enzyme activities**

**4a) Trehalose 6-phosphate phosphatase activity**

By monitoring release of inorganic phosphate (Pi) from crude extract using the malachite-green reagent (Sigma-Aldrich, MA, USA) to assess the enzymatic activity of trehalose 6-phosphate phosphatase (TPP). The assay was conducted following the modified protocol of Klutts et al. (2003). The reaction was performed in a total volume of 100 µL with a final concentration of 1 mM trehalose 6-phosphate (Sigma-Aldrich, MA, USA), 2 mM MgCl₂ (Hi-Media, MS, India), 50 mM citrate-phosphate buffer (pH 4), and 35 μg of enzyme extract. The reaction mixture was then incubated at 57°C for 35 minutes to allow for hydrolysis. The Klutts procedure was followed, wherein the reaction was first quenched by the addition of two volumes of malachite green 0.15% filtered, 1% ammonium molybdate (Hi-Media, MS, India), and 12.5% (v/v) concentrated HCl (Thomas Baker, MS, India). After spectrophotometric measurements of the absorbance at 630 nm, sediment was separated and color terbium titration was held. Color was developed for 5-7 minutes. It can be claimed that this composition effectively measures Pi release.

**4b) Trehalase activity**

The α,α-trehalase activity was performend on respective assays by calculating the percentage of residual glucose that was hydrolyzed from α,α-trehalose using the Dinitrosalicylic acid (DNSA) reagent. A premix containing buffer and equal volumes of crude sample extracts from the control and treatment groups were prepared. To this rection, 150 μL of trehalose i.e. 0.25% was added. The reaction was allowed to hydrolyze the enzyme for 15 minutes at 37oC. The reaction was then stopped with addition of 500 μL of DNSA reagent. The mixture was then placed in a boiling water bath for 5 minutes in order to enhance the detection of glucose through colorimetry. The quantitative measurement of glucose released during the trehalose hydrolysis was single at 540 nanometres. The absorbance of the measurement was retrieved at 540 nm. Under the defined conditions of the assay, one unit of trehalase activity was defined as the amount of enzyme liberated of 1μM of glucose per minute.

**Supplementary method 5**:

LC-MS Method- To extract metabolite 80-100mg of finely crushed sample powder was mixed with **500 µL of 80% MS-grade methanol**, to ensures efficient metabolite extraction. Then this mixture vortexed for 15-20 min at room temperature to enhance homoginization, then proceed for 20 minutes sonication to disrupt cellular content followed by centrifugation at 18000 × g for 10-20 minutes to get clear supernatant. To stabilizes metabolites and prevents further degradation supernatant stored in -80°C deep freezer. The next day, samples were again centrifuged at high speed for 10-20 min to remove of residual particales and supernatant filtered through a 0.2 µm syringe filter to remove **micro-particales** that may interfere with detection. The LC method started with 2% B for the first 0.3 min that is the conditions the column for metabolite separation and for increase compound resolution solvent strength increased to 30% in the next 2 min. The B% was increased from 30 to 45% till 5 min and further increased to 98% to 7 min at which it was held for the next 50sec.The column was equilibrated to the initial ratio of solvents (98% A: 2% B) in the last 2 min to restores initial conditions for the next run.The MS data from two or three independent biological replicates and two technical replicates each were acquired in 5 GHz extended dynamic range.
