## Supplementary material for "Trehalose Transport Dynamics Underpin a Metabolic Trade-Off between Exogenous Uptake and Endogenous Synthesis in Lepidopteran Insects": Supplementary File_S5-Results-Ha Only.docx

Figure S1: Average body weight of *Helicoverpa armigera* larvae upon different concentrations of trehalose feeding

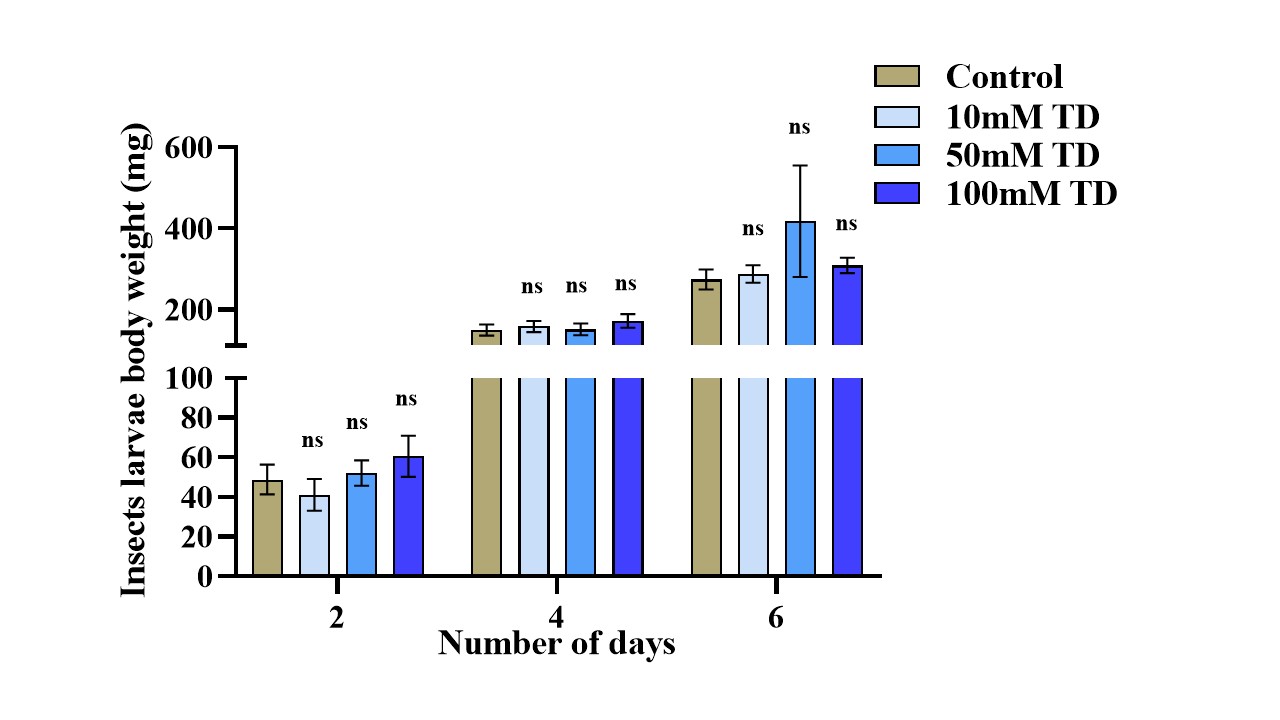

Table S2: Nutritional indices of *Helicoverpa armigera* larvae upon different concentrations of trehalose feeding

|  | Nutritional indices (Mean±SE) | | | | | | | | |
| --- | --- | --- | --- | --- | --- | --- | --- | --- | --- |
| Assaysets | ECI |  |  | ECD |  |  | ADI |  |  |
|  | Day-3 | Day-5 | Day-7 | Day-3 | Day-5 | Day-7 | Day-3 | Day-5 | Day-7 |
| Control | 17.7323±2.868 | 52.099±3.6601 | 75.59±2.576 | 20.73±3.39 | 69.87±4.12 | 1.22±4.66 | 89.02±2.73 | 75.255±3.318 | 24.40±2.576 |
| 10mM | 13.75±2.96 | 47.73±3.15 | 73.27±3.632 | 15.41±3.23 | 65.52±4.325 | 111.11±5.44 | 92.30±2.31 | 75.16±3.516 | 67.854±3.271 |
| 50mM | 18.879±2.99 | 52.10±3.669 | 68.77±4.302 | 22.19±3.480 | 75.25±4.586 | 116.73±5.160 | 88.025±2.516 | 72.155±4.080 | 58.61±2.710 |
| 100mM | 21.33±3.029 | 51.69±3.682 | 82.38±3.4939 | 26.86±3.694 | 80.461±5.147 | 198.095±13.53 | 82.82±2.811 | 66.84±3.256 | 53.98±4.05 |

Figure S3: Expression dynamics of *HaSTs* upon trehalose feeding

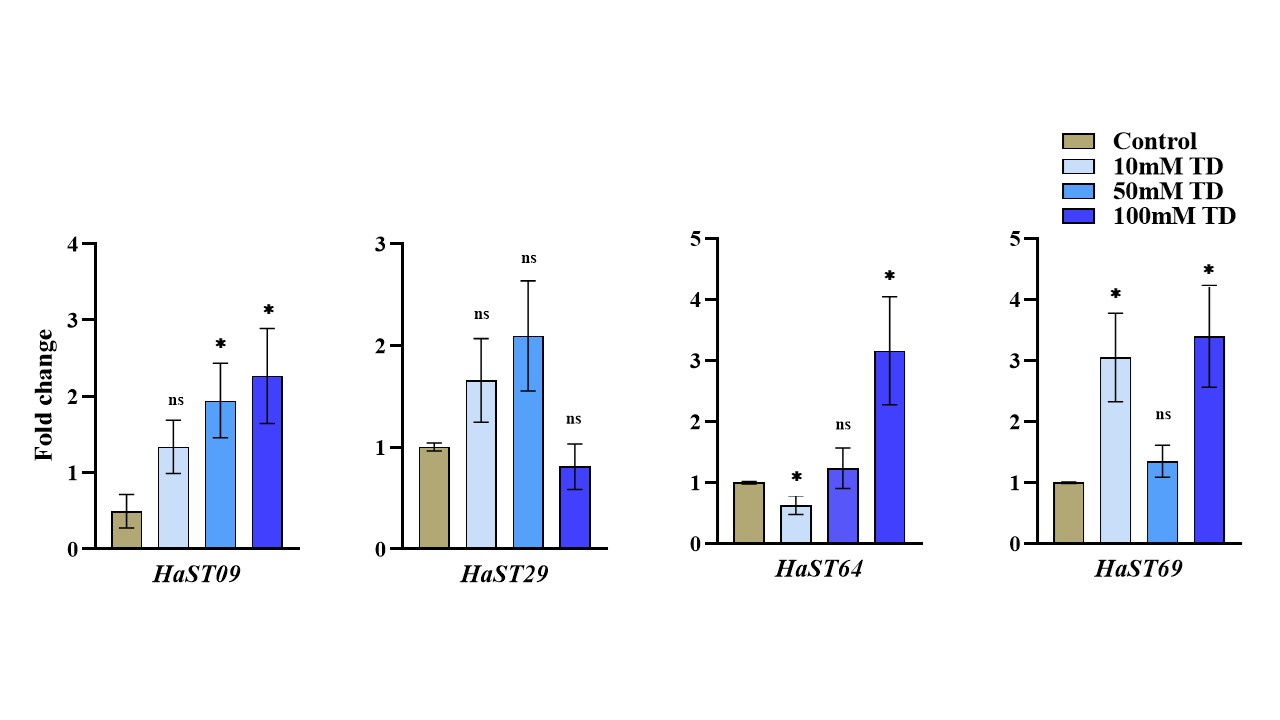

Figure S4: Expression dynamics of *HaSTs* upon 50mM trehalose feeding

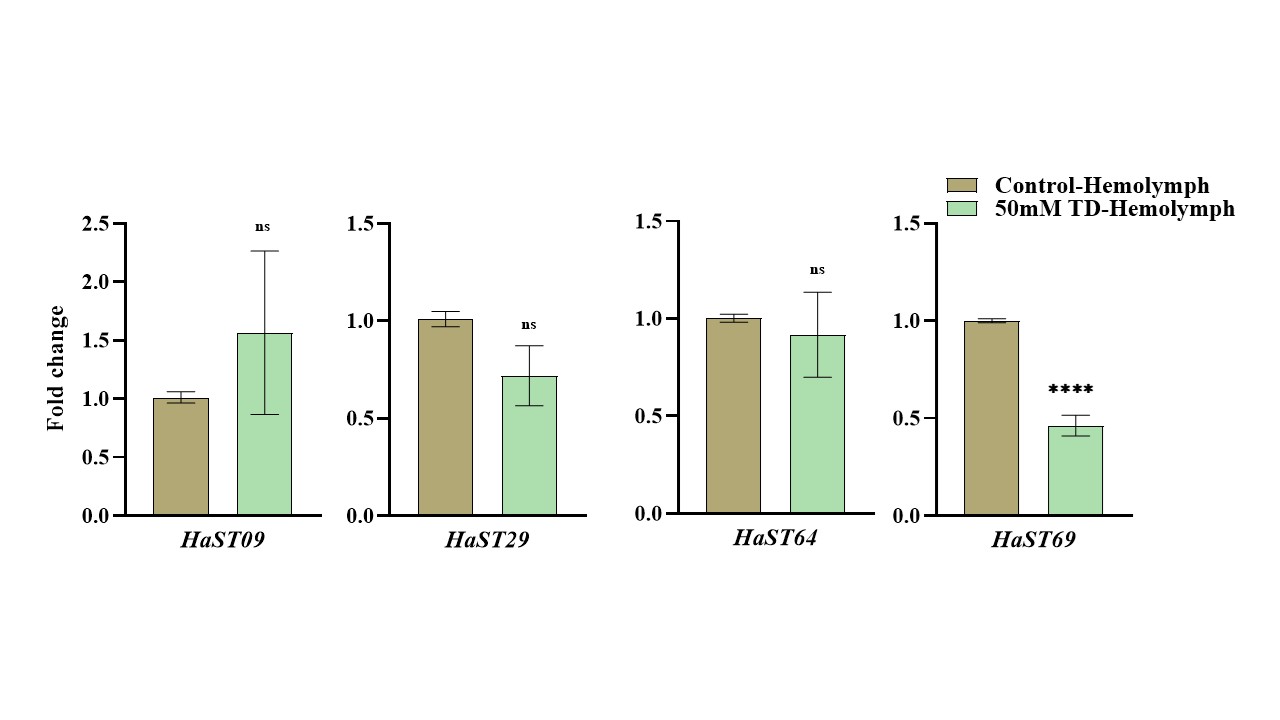

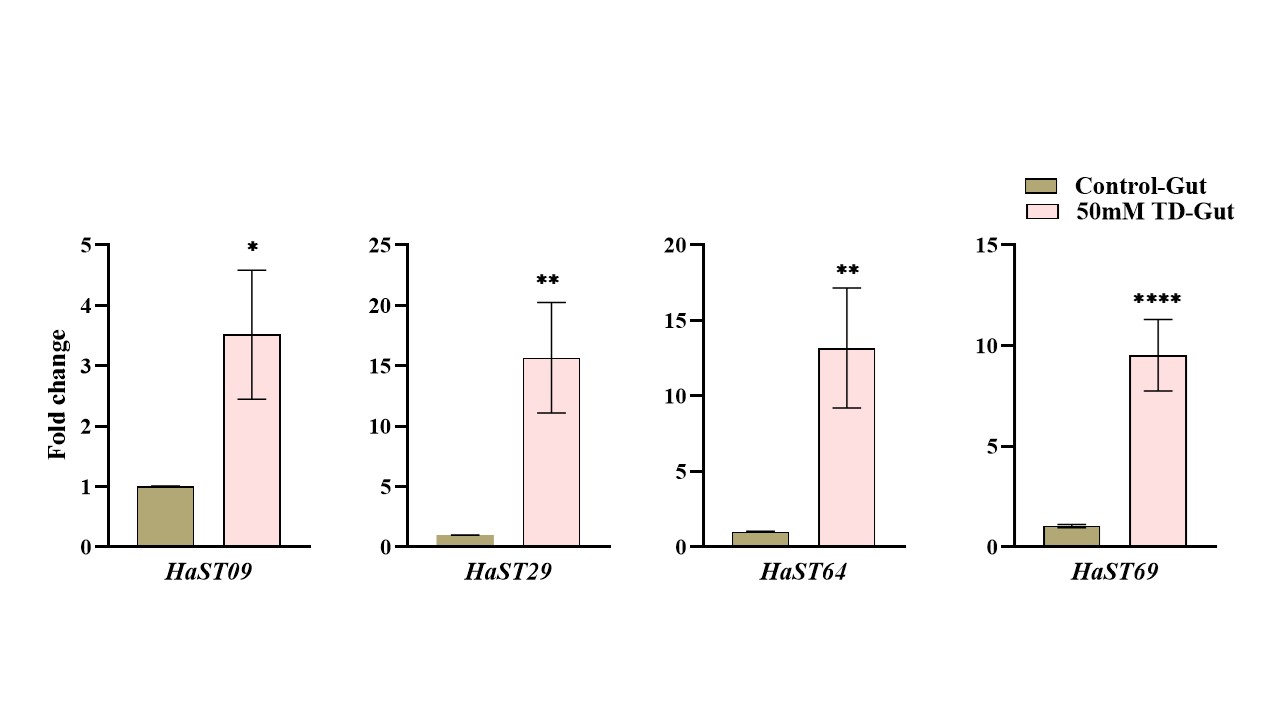

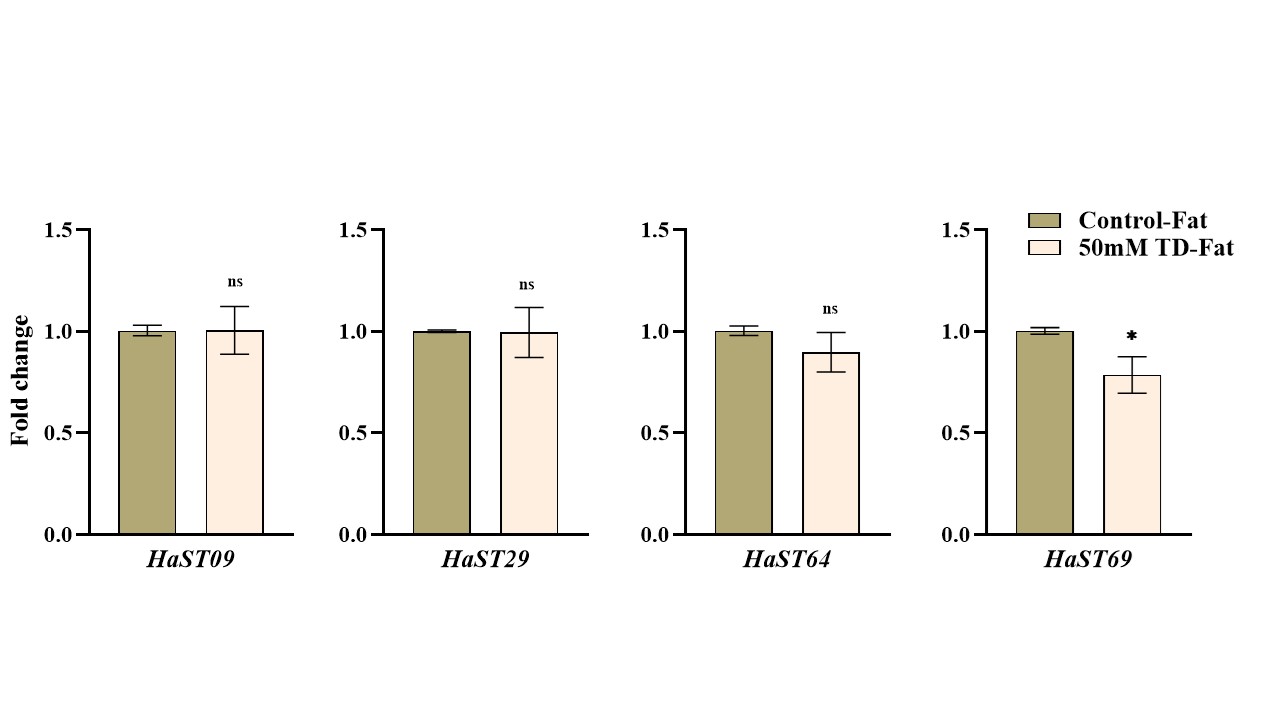

Figure S5: Trehalose levels quantification in hemolymph upon 50Mm trehalose feeding using LC-MS/MS

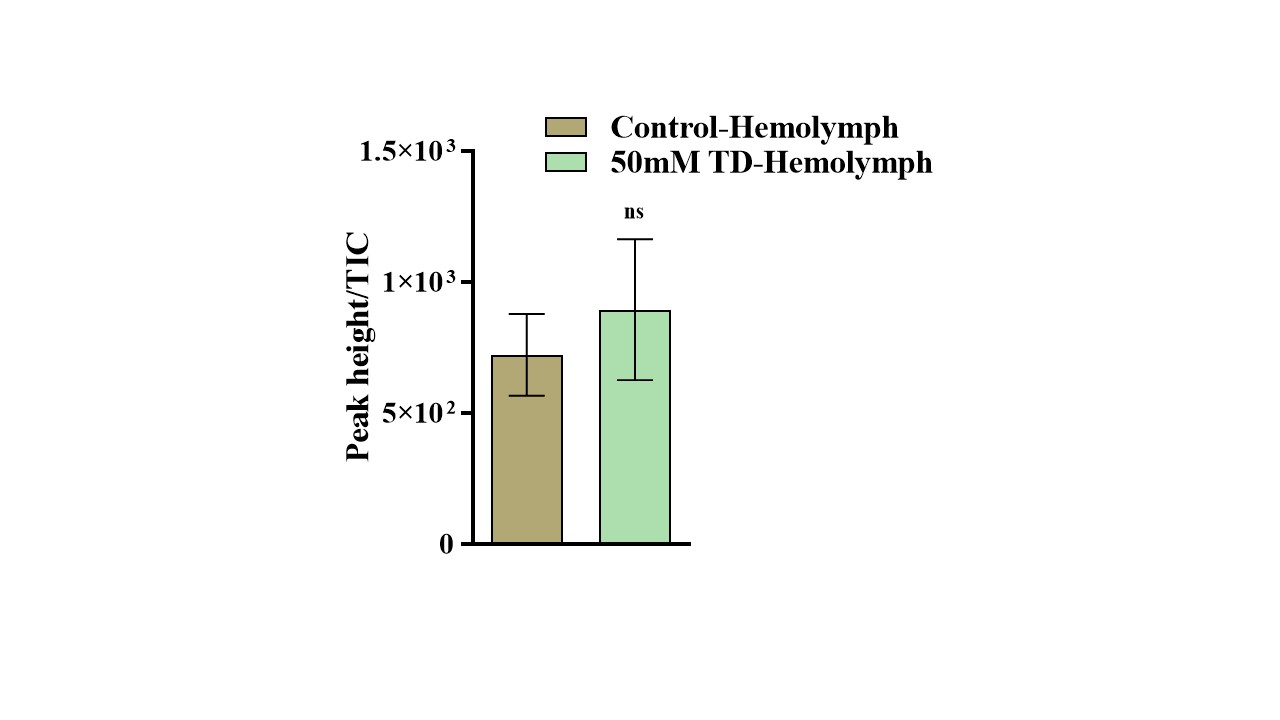

Figure S6: *dsHaST46* clone confirmation in L4440 vector by restriction digestion

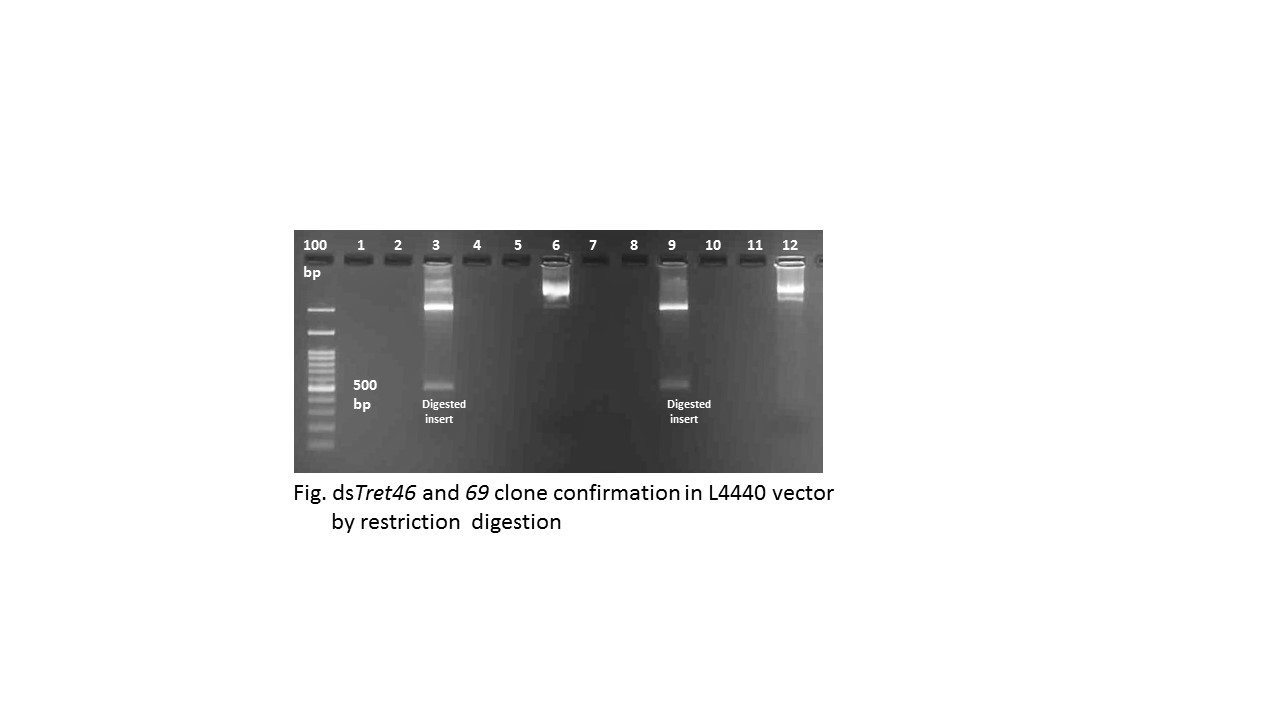

Figure S7: Confirmation of ds*HaST46* production via RNA isolation and visualization on 1.2% agarose gel (ds*ST46*expressed band seen around 630bp). 400bp amplicon plus flaking regions from pGEM-T easy vector and L4440 vector.

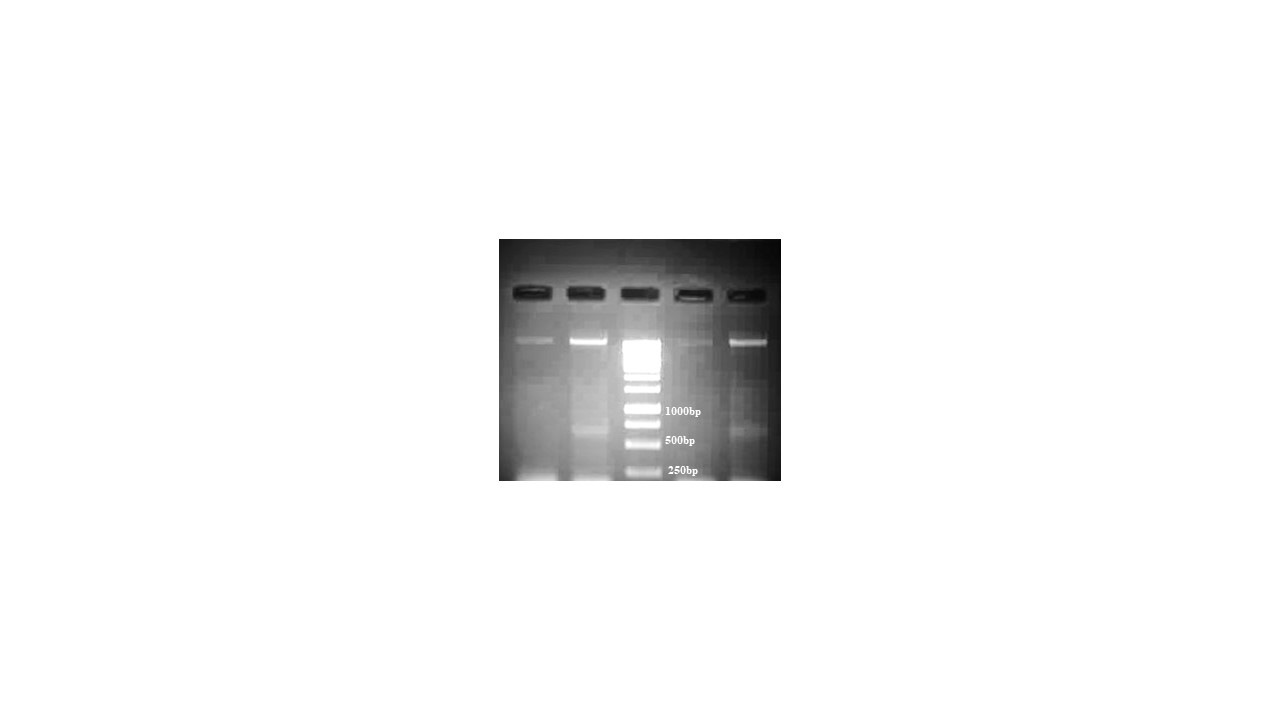

***dsHaST46***

**L4440**

Figure S8: Average body weight of *H. armigera* larvae upon empty vector (L4440) and *dsHaST46*-treated insects, either fed on an artificial diet or an artificial diet supplemented with 50 mM trehalose

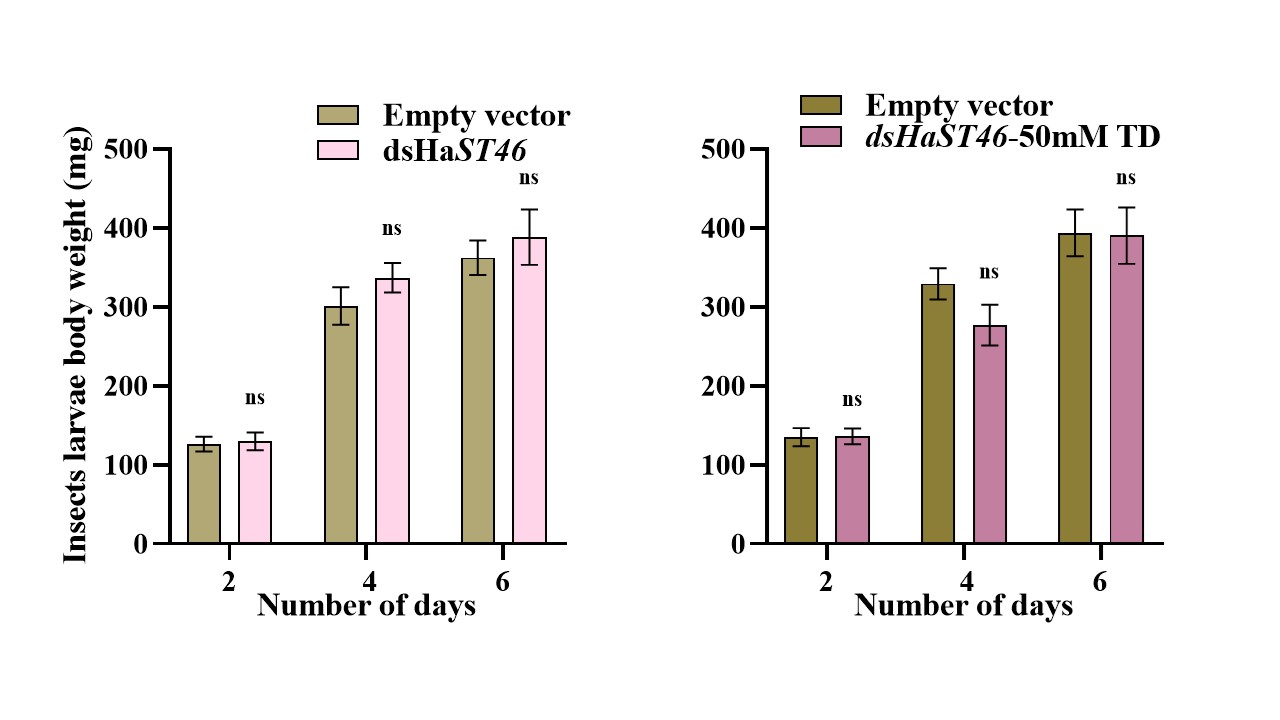

Table S9: Effect of ds*HaST46* ingestion on *H. armigera* feeding and dietary utilization

|  | Nutritional indices (Mean±SE) | | | | | | | | |
| --- | --- | --- | --- | --- | --- | --- | --- | --- | --- |
| Assay sets | ECI |  |  | ECD |  |  | ADI |  |  |
|  | Day-3 | Day-5 | Day-7 | Day-3 | Day-5 | Day-7 | Day-3 | Day-5 | Day-7 |
| Empty vector | 35.17±2.313 | 47.29±3.00 | 69.16±3.24 | 45.46±2.87 | 80.090±3.586 | 147.26±4.642 | 78.22±2.97 | 60.164±3.549 | 47.04±1.697 |
| ds*HaST46* | 38.22±2.696 | 46.60±3.105 | 72.143±3.273 | 50.76±3.39 | 77.48±3.8803 | 147.44±3.280 | 76.66±3.090 | 60.85±3.093 | 49.22±2.9685 |

Figure S10: Insect phenotype in *H.armigera* larvae fed on bacterial-expressed dsRNA

a

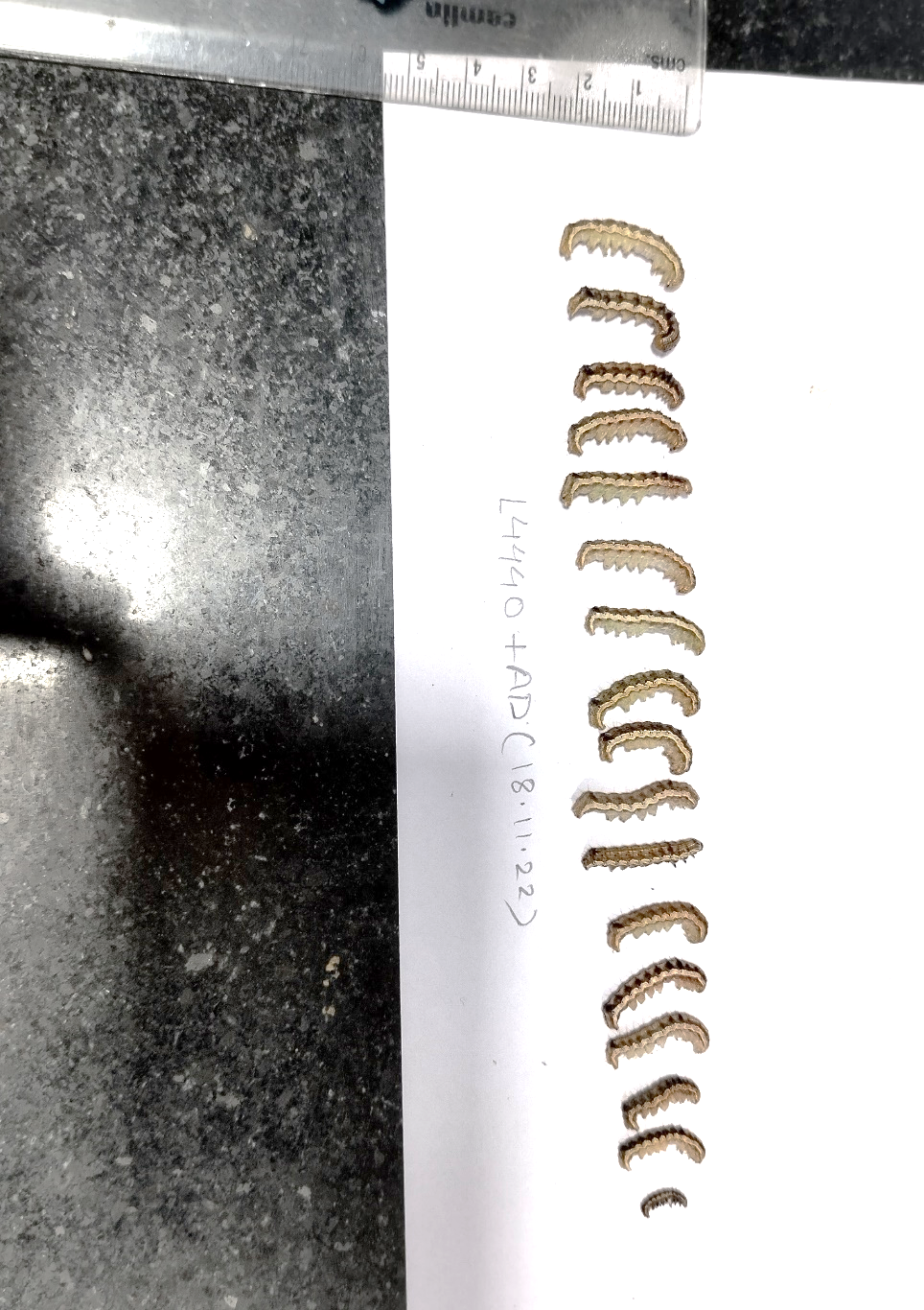

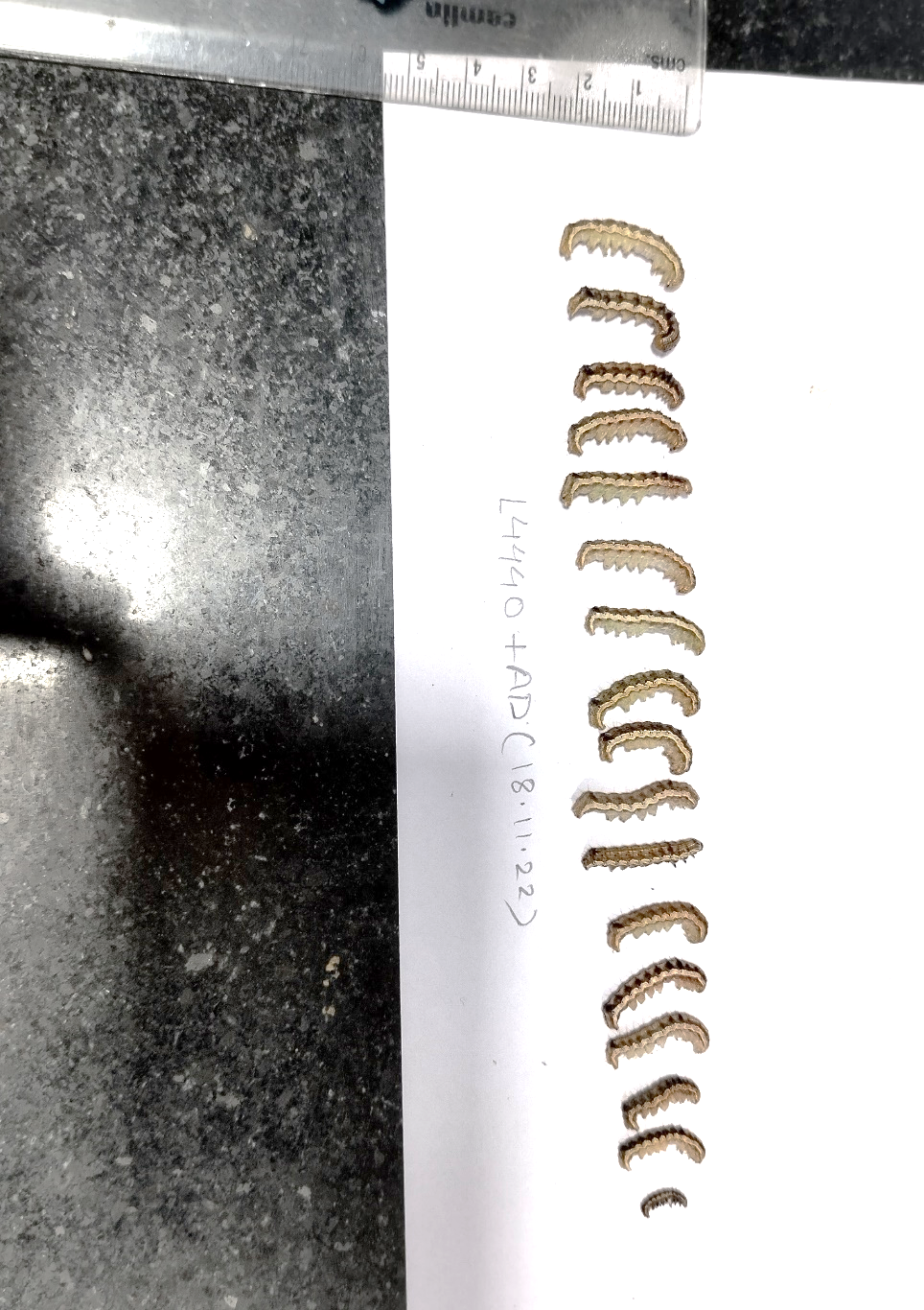

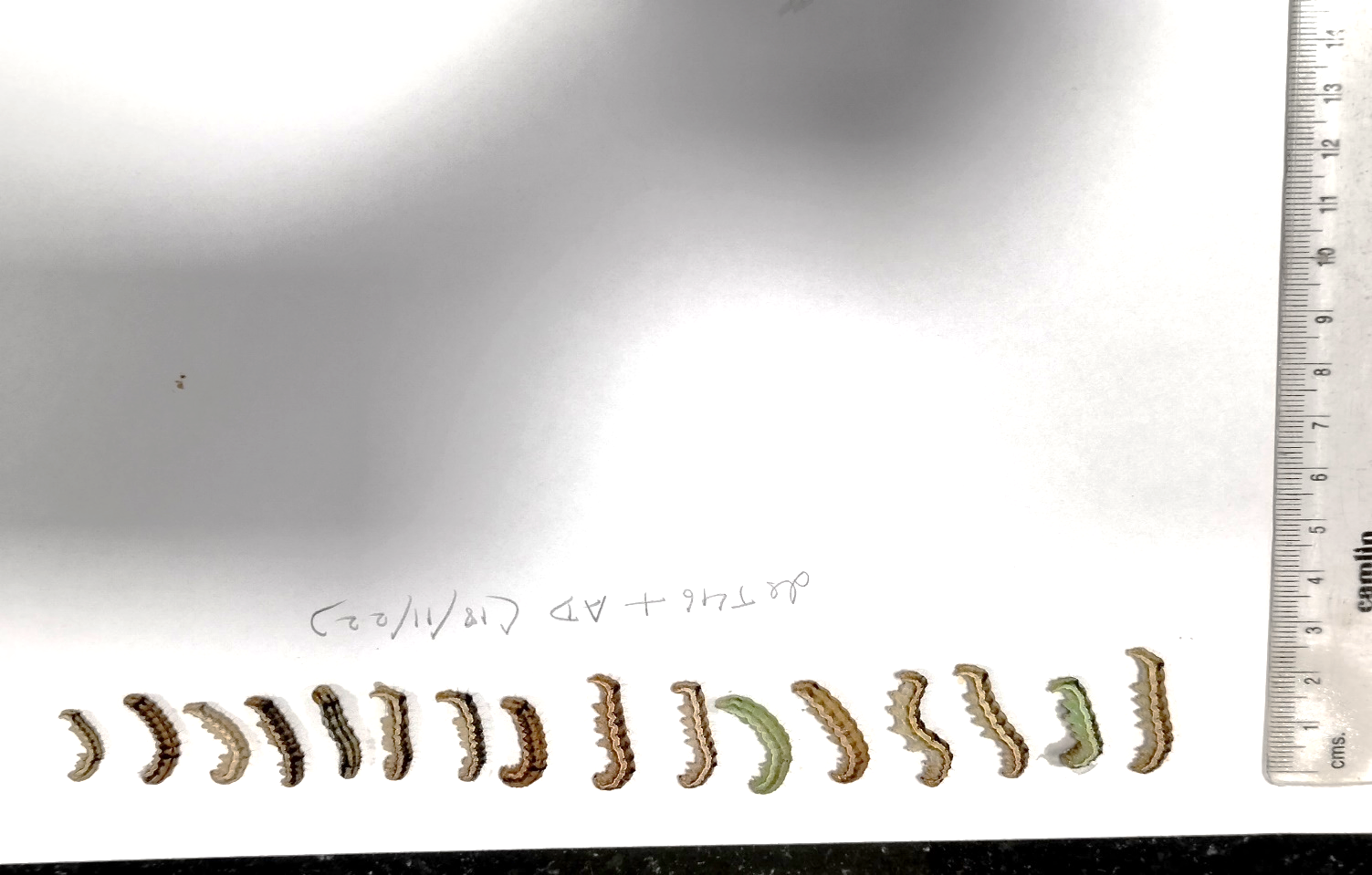

b

c

d

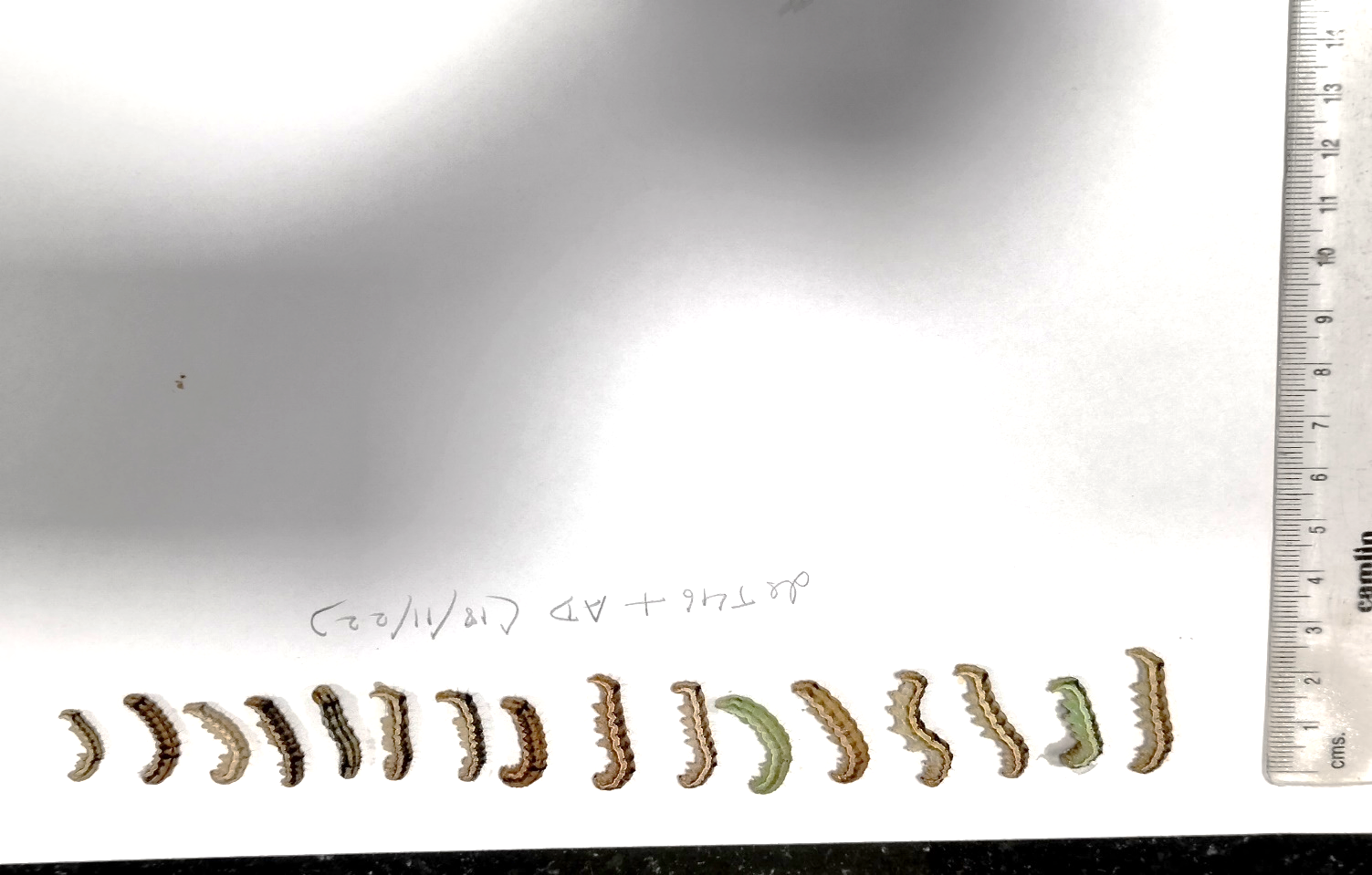

Figure S11: Insect phenotype in *H.armigera* larvae fed on bacterial-expressed dsRNA supplemented with 50 mM trehalose (a) Insect fed on bacterial-expressed dsRNA of L44440 vector (b) Insects fed on bacterial-expressed dsRNA of HaST46(c) Insect fed on bacterial-expressed dsRNA of L44440 vector supplemented with 50 mM trehalose (d) Insects fed on bacterial-expressed dsRNA of HaST46 supplemented with 50 mM trehalose

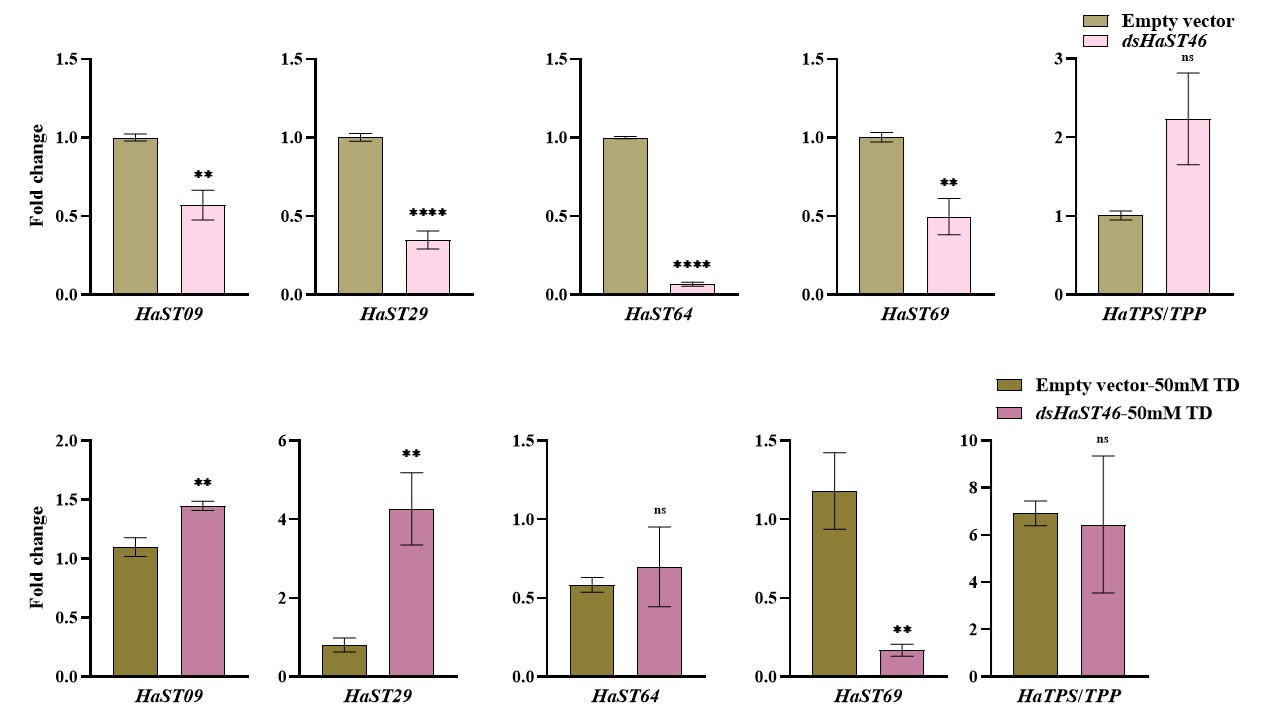

b

a

Figure S12 a-b: qRT-PCR analysis of *HaSTs* and *HaTPS/TPP* expression in whole-body samples from control insects (empty L4440 vector) and *dsHaST46*-treated insects, either fed on an artificial diet or an artificial diet supplemented with 50 mM trehalose

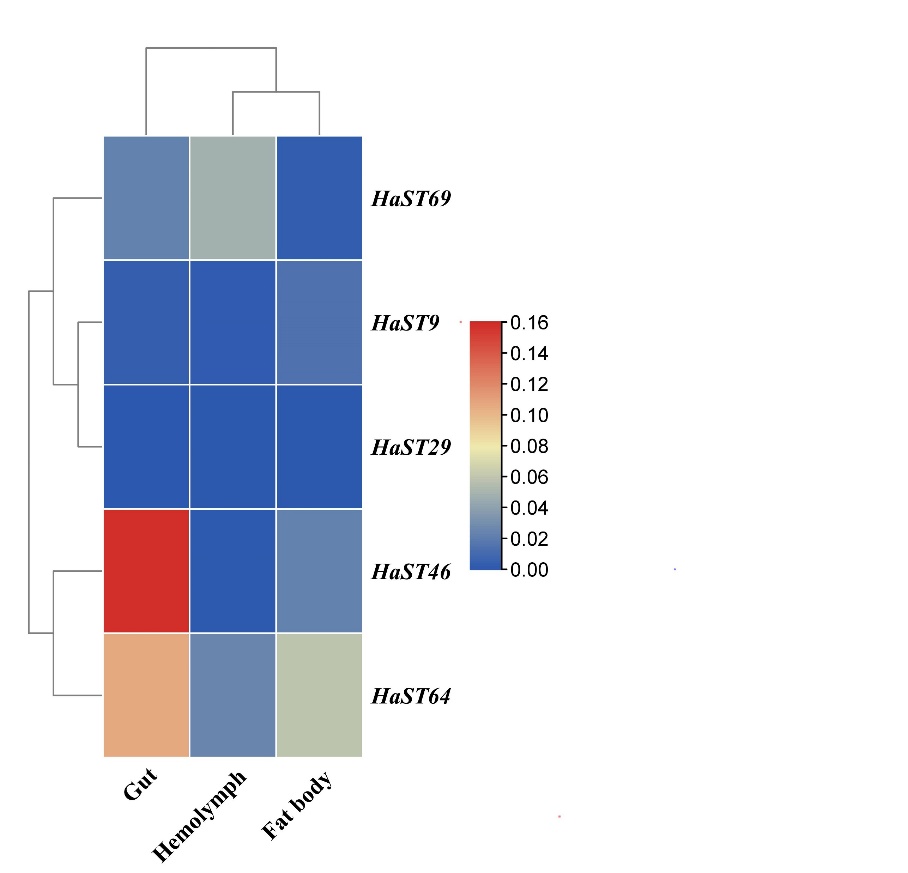
Figure S13: DGE analysis of *HaSTs* across tissues
